# Real-Time Linear Prediction of Simultaneous and Independent Movements of Two Finger Groups Using an Intracortical Brain-Machine Interface

**DOI:** 10.1101/2020.10.27.357228

**Authors:** Samuel R. Nason, Matthew J. Mender, Alex K. Vaskov, Matthew S. Willsey, Parag G. Patil, Cynthia A. Chestek

## Abstract

Modern brain-machine interfaces can return function to people with paralysis, but current hand neural prostheses are unable to reproduce control of individuated finger movements. Here, for the first time, we present a real-time, high-speed, linear brain-machine interface in nonhuman primates that utilizes intracortical neural signals to bridge this gap. We created a novel task that systematically individuates two finger groups, the index finger and the middle-ring-small fingers combined, presenting separate targets for each group. During online brain control, the ReFIT Kalman filter demonstrated the capability of individuating movements of each finger group with high performance, enabling a nonhuman primate to acquire two targets simultaneously at 1.95 targets per second, resulting in an average information throughput of 2.1 bits per second. To understand this result, we performed single unit tuning analyses. Cortical neurons were active for movements of an individual finger group, combined movements of both finger groups, or both. Linear combinations of neural activity representing individual finger group movements predicted the neural activity during combined finger group movements with high accuracy, and vice versa. Hence, a linear model was able to explain how cortical neurons encode information about multiple dimensions of movement simultaneously. Additionally, training ridge regressing decoders with independent component movements was sufficient to predict untrained higher-complexity movements. Our results suggest that linear decoders for brain-machine interfaces may be sufficient to execute high-dimensional tasks with the performance levels required for naturalistic neural prostheses.

## INTRODUCTION

Neural prostheses have the potential to return independence to many people with neurological disorders or injuries. In human clinical trials, laboratories have restored use of computers, self-feeding, and prosthetic hands using implants to translate electrophysiological signals into user intent (Ajiboye et al., 2017; Memberg et al., 2014; Nuyujukian et al., 2016; Pandarinath et al., 2017; Wodlinger et al., 2015). Of greatest interest to people with cervical-level spinal cord injury is the return of hand and arm function (Anderson, 2004). Although this has motivated many groups to study neural prostheses for hand control, only a few have been translated to use with people and none have been translated to full-time use outside of the laboratory. Functional electrical stimulation provides an avenue for outputting the intentions of the user to their natural limbs (Kilgore et al., 1989, 2008; Memberg et al., 2014; Smith et al., 2005), with commercial solutions having already existed such as the FreeHand System. Unfortunately, they typically rely on some external motion or myoelectric commands from residual functional muscles, which require learning and are generally unnatural to use.

This has driven many groups to use brain-machine interfaces to extract hand prosthesis control signals from a more natural source. In humans, various studies have attempted to characterize the relationship between finger movements and electrocorticography activity (Chestek et al., 2013; Hotson et al., 2016; Kubánek et al., 2009). However, that relationship was insufficiently strong to enable quick classifications or fully dexterous continuous movements. Some groups have used intracortical microelectrodes in monkeys to record activity patterns on the order of single neurons to investigate a more concrete connection between them and finger behaviors (Baker et al., 2009; Mollazadeh et al., 2011). These studies suggest such a relationship is stronger than with electrocorticography, but classification of which finger is moving by itself does not provide enough understanding of that relationship to predict the quality of continuous control.

The ability to precisely control the positions of all individual fingers is a key characteristic of dexterous hand use in primates. As such, many groups have attempted to relate many, if not all 27, of the degrees of freedom (DoFs) within the hand to neural activity during reach-to-grasp tasks offline (Aggarwal et al., 2013; Bansal et al., 2011; Okorokova et al., 2020; Vargas-Irwin et al., 2010). Although these studies showed very high offline correlation between firing rates and behaviors for many of those DoFs, most, if not all, of the DoFs presented showed highly correlated trajectories and may not have truly been independent. This makes it unclear how well this neural activity corresponds to those DoFs individually or if all the DoFs are moving so similarly that anything with a similar time course will correlate well. Further, without evaluating online control of individual DoFs, the applicability to intuitive and naturalistic neural prostheses is uncertain.

Generally, the algorithms for online control of neural prostheses assume linear relationships between primary motor cortex neural activity and either the position and velocity or expected muscle activations of prosthetic movements. Variants of Kalman filters (Ajiboye et al., 2017; Gilja et al., 2012; Malik et al., 2011; Wu et al., 2004), ridge regressions (Collinger et al., 2013; Mulliken et al., 2008; Wodlinger et al., 2015), and Wiener filters (Ethier et al., 2012; Koyama et al., 2010; Sachs et al., 2015) have been used to control arms, hands, and fingers online. Linear online decoders are promising candidates for an out-of-laboratory clinical neural prosthesis due to their computational simplicity and high prediction performance. However, with limited quantities of recording electrodes, covariances between nearby neural signals, and increasing numbers of DoFs required for finger control, linear decoders may be unable to accommodate multiple independent degrees of freedom.

It has been noted by several groups that the same neurons can covary with substantially different behaviors, which could make the prediction of finger movements particularly difficult. For example, primary motor cortex can simultaneously encode information about upper extremities, fingers, and speech, independent of body laterality (Cross et al., 2020; Diedrichsen et al., 2013; Heming et al., 2019; Jorge et al., 2020; Stavisky et al., 2019, 2020; Willett et al., 2020). As tasks increase in complexity, linear models may be unable to discriminate between neural states without sampling greater quantities of relevant neurons. Therefore, it is valuable to characterize the limits of linear models in discriminating neural states with truly simultaneous movement of independent DoFs.

Here, we show for the first time fine, independent, and simultaneous online control of two systematically individuated groups of fingers within one hand to acquire two targets, one each for the index finger and the middle-ring-small (MRS) fingers, using linear Kalman filters and an intracortical brain-machine interface in nonhuman primates. With intention-based retraining of the Kalman filters, we find that online brain control improves significantly. Then, we find that the magnitude of individual neural activations to particular movements, whether they correspond to movements of one group or combined movements of both groups, can be well predicted by the weighted sums of the most similar movements. This suggests that neural representations of continuous finger movements are linearly related. Finally, we train offline ridge regression decoders on either individual or combined finger movements and test on the untrained movements to show that population-level neural activations for our two-dimensional finger task are mostly consistent between individual and combined finger group movements.

## RESULTS

### Linear Two-Finger Decoding in Real-Time

We first sought to validate that linear decoder models could individuate two systematically separated finger dimensions moving independently and often simultaneously throughout their entire ranges of motion. To do so, we trained two adult male able-bodied rhesus macaques, monkeys N and W, to perform a two-target two-finger acquisition task by pushing two doors on a manipulandum, one door for each finger group (Vaskov et al., 2018). This task is illustrated in Figure 1A. Although the monkeys were presented with two targets, they can be visualized as one target in a two-dimensional space of % index flexion versus % MRS flexion, as shown in Figure 1B. Three magnitudes were pseudo-randomly presented for each posture (+20%, +30%, or +40% from rest), resulting in 19 total target combinations without the MI and IM posture styles (see methods). During the task, we synchronously recorded neural activity using 96 channels of implanted Utah silicon microelectrode arrays (Blackrock Microsystems, Salt Lake City, UT, USA) from the hand area of primary motor cortex in each monkey (implant photographs in Figure 1D). Each experimental day, we collected a training dataset in which the monkey controlled the virtual hand using the manipulandum while synchronously recording 300-1,000Hz spiking band power (SBP). We have previously shown that SBP is well correlated with the firing rate of the largest amplitude single unit or units on an electrode and typically results in higher decoding performance than threshold crossing rate (Nason et al., 2020).Then, we trained a Kalman filter as detailed in the methods to predict fingertip velocities in real-time and tested it in closed-loop by actuating the virtual hand according to the predictions.

**Figure 1.**
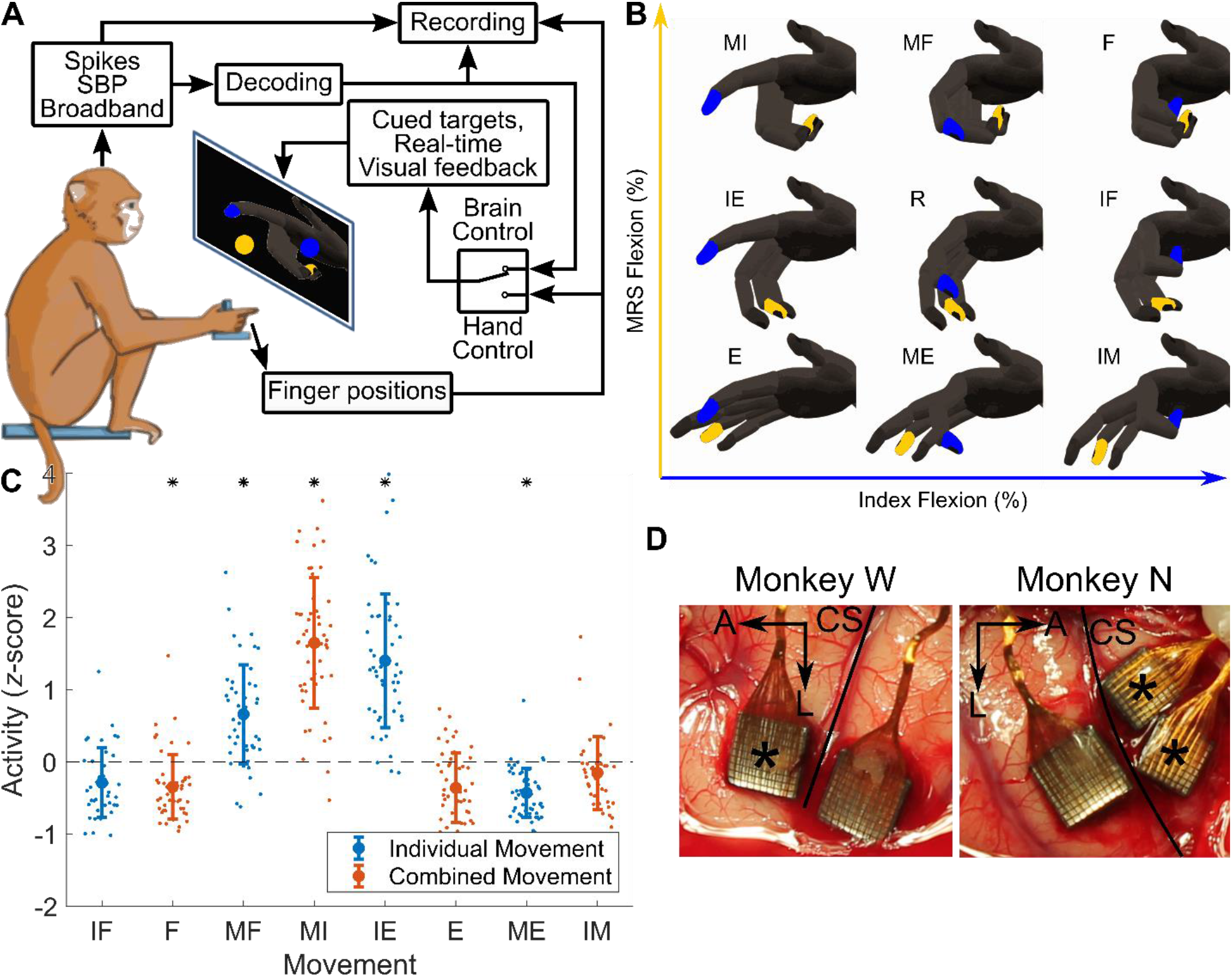
Experimental and system description. **(A)** Flow diagram of information through the experimental system. Positions of the index and middle/ring/small (MRS) finger groups are measured synchronously with the neural activity, namely spikes, spiking band power (SBP), and broadband (0.1Hz-7.5kHz sampled at 30kSps). The finger position measurements or the decoded finger positions are used to actuate the virtual hand, depending on the stage of the experiment. **(B)** Two-dimensional space for visualizing the hand movements. I - index finger group, M - MRS finger group, F - flexion, E - extension, IM - index flexion with MRS extension, MI - MRS flexion with index extension, R - rest. **(C)** An example tuning curve from monkey N illustrating an SBP channel tuned to index extension movements. Asterisks indicate significant difference from the average activity across the experiment (two-sided two-sample Kolmogorov-Smirnov test, *p* < 0.001, corrected for false discovery rate). **(D)** Photographs of monkey W’s and monkey N’s intracortical Utah microelectrode array implants. Monkey W’s implants were in his left hemisphere and monkey N’s implants were in his right hemisphere. * indicates arrays used in this study. A - anterior, L - lateral, CS - central sulcus.

We found that the monkeys could successfully control the movements of both finger groups independently in real-time using a linear Kalman filter. Figure 2A and B show representative predicted finger traces from the SBP Kalman filter for monkeys N and W, respectively, that show a wide variety of movements. These traces demonstrate that they could individuate the two fingers independently with smooth and controlled effort using the brain-machine interface. Figure 2C shows the closed-loop statistics for the two-finger SBP Kalman filter. Regarding acquisition time, monkey N acquired targets at an average 1.8s using the decoder, greater than the average 0.87s acquisition time when controlling the virtual hand using the manipulandum. The other metric we investigated is information throughput, which is a measure of task difficulty balanced by the time taken to complete it, calculated via Fitts’ Law as detailed in the methods. Monkey N achieved an average 1.7bps with the Kalman filter, which was lower than the average 2.5bps achieved with the manipulandum. In contrast, even with only 30 minimally modulated SBP channels and no modulated sorted units, monkey W was able to complete 78% of trials. Not surprisingly from the degraded neural signals, the average 11s acquisition time and average 0.48bps bit rate were substantially degraded from manipulandum control (1.2s acquisition time and 2.3bps throughput). Nonetheless, the traces and statistics demonstrate individuation in closed-loop brain control. For completeness, Supplementary Figure 1 illustrates the same for threshold crossing rate rather than SBP, but only for monkey N, as monkey W could not complete the task with any success using the threshold crossing rate Kalman filter. This was expected, as we have shown previously that SBP’s small passband enables the extraction of the firing rates of single units at lower signal-to-noise ratios (SNRs) than high bandwidth methods (Nason et al., 2020).

**Figure 2.**
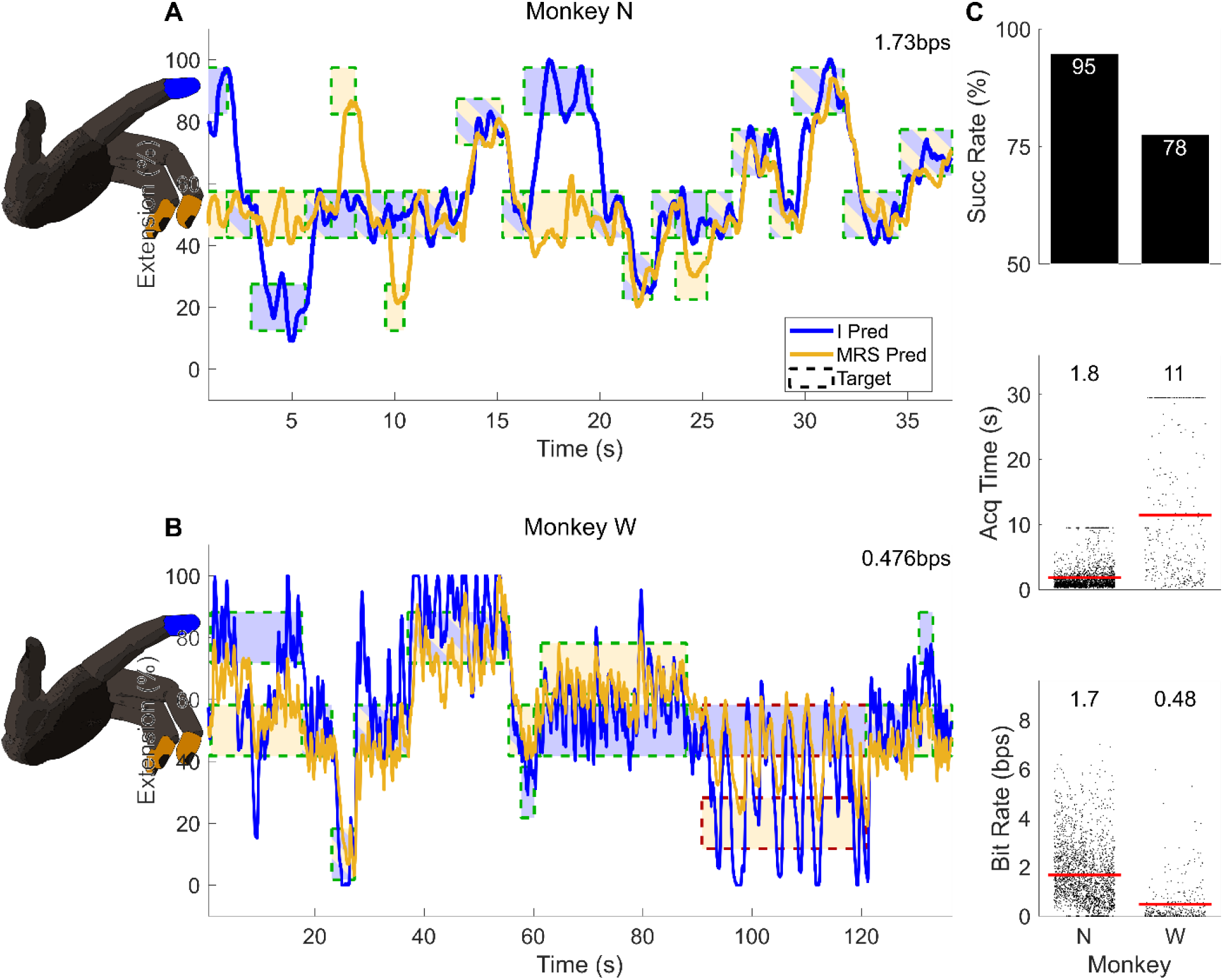
Two-finger closed-loop Kalman filter decodes using spiking band power (SBP). **(A, B)** Example closed-loop prediction traces from monkeys N and W using the standard Kalman filter, respectively. Targets are represented by the dashed boxes, internally colored to indicate the targeted finger with a border color representing whether the trial was acquired successfully (green, red if not). “I” means the index finger group and “MRS” means the middle/ring/small finger group. The mean bit rate of the trials displayed in each window is presented at the top right. **(C)** Statistics for all closed-loop two-finger Kalman filter decodes for monkeys N (left) and W (right). The red lines indicate the means, which are numerically displayed above each set of data. The statistic for each trial is represented by one dot in each plot. “Succ Rate” means the percentage of total trials that were successfully acquired in time, and “Acq Time” means target acquisition time.

Out of interest for applications to brain-machine interfaces, we attempted to maximize closed-loop, two-finger decoding performance using the state-of-the-art recalibrated feedback intention-trained (ReFIT) Kalman filter (RFKF; Gilja et al., 2012; Vaskov et al., 2018). The RFKF training procedure occurs following the monkey’s usage of the original Kalman filter. It assumes that the neurons controlling the Kalman filter’s predictions represented the intention of the monkey to optimally bring the fingers to the targets, regardless of the directions of the predictions. Then after reorienting predictions to match the presumed intentions of the monkey, the linear model is retrained. As discussed in the methods, the two interpretations of our finger task result in two frameworks for retraining: rotation of the net velocity in two-dimensional finger space to back-calculate each finger’s intended velocity (similar to the original ReFIT method) or independent negation of each finger’s velocity if it is moving away from the target. The statistics comparing these two methods and a third combining both are shown in Figure 3C. Ultimately, there was no statistical difference between any pair of recalibration methods (*p* > 0.001, two-tailed two-sample *t*-test).

**Figure 3.**
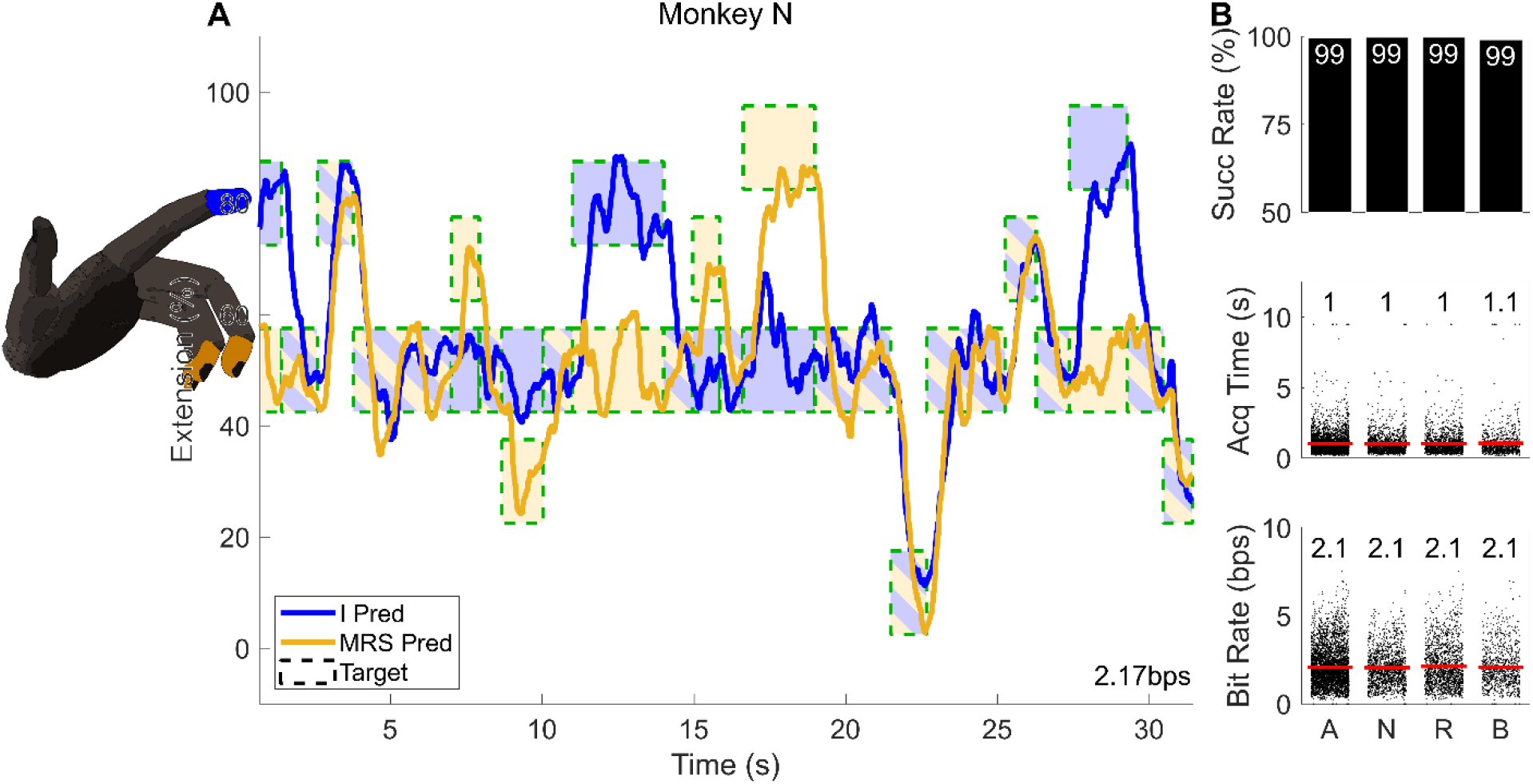
Two-finger closed-loop ReFIT Kalman filter decodes. (**A**) Example closed-loop prediction traces from monkey N using the ReFIT Kalman filter. Targets are represented by the dashed boxes, internally colored to indicate the targeted finger. “I” means the index finger group and “MRS” means the middle/ring/small finger group. The mean bit rate of the trials displayed is presented at the bottom right. (**B**) Statistics for all closed-loop two-finger ReFIT Kalman filter decodes. The red lines indicate the means, which are numerically displayed above each set of data. The statistic for each trial is represented by one dot in each plot. “Succ Rate” means the percentage of total trials that were successfully acquired in time, and “Acq Time” means target acquisition time. (**C**) Statistics of each type of prediction reorientation for ReFIT training. “N” means velocities for each finger were negated if not pointing to that finger’s target, “R” means the velocities were rotated in the two-dimensional finger space towards the target, and “B” means both reorientations were used by concatenating them and repeating the neural activity.

Overall, the RFKF made a substantial improvement in decode performance over the standard Kalman filter. Figure 3A shows closed-loop prediction traces from monkey N using the SBP RFKF. In comparison to the Kalman filter, ReFIT Kalman filters significantly improve prediction performance (1.0s versus 1.8s acquisition time and 2.1bps versus 1.7bps throughput for monkey N, *p* < 0.001, two-tailed two-sample *t*-test). Additionally, the predictions are less oscillatory when attempting to stop on a target, and when it is oscillating, the amplitude is generally smaller than the standard SBP Kalman filter, aligning with what was previously reported (Gilja et al., 2012, 2015; Vaskov et al., 2018). To showcase this, Supplementary Video 1 presents monkey N’s usage of the SBP RFKF in real-time with a general comparison to manipulandum control.

### Cortical Neurons Show Specificity to Individual Contractions

We found it surprising that such a complex hand task that humans would likely find difficult to perform could be captured so well by a linear decoder. To look at whether these linear relationships hold within individual neurons, we constructed tuning curves of individual units across all eight finger postures at just the +30% magnitude. On one exclusive and representative day for each monkey, they performed the two-finger task in manipulandum control for at least 30 continuous minutes while threshold crossings, SBP, and broadband activity were recorded synchronously. The broadband activity was spike sorted using Offline Sorter (Plexon, Dallas, TX, USA) to extract the firing rates belonging to single units. 386 trials for monkey N and 125 for monkey W were processed. We calculated the mean firing rate for each type of movement and plotted tuning curves. Illustrative tuning curves are displayed in the left plots of Figure 5A and C. Initially, we characterized the tuning preferences of our 60 units, which are summarized in Table 1 and Table 2. First, in Table 1 regarding specificity of neural activation to flexion and extension, we found 20 of the 60 units showed specificity to one muscle group but not the other, suggesting cortical representation of distinct muscle groups. Second, in Table 2 regarding specificity to individuated finger group movements, we found that 18 of the 60 units showed tuning to only one finger group. Among the units that showed specificity to one group, there were 50% more units tuned to movements of the MRS finger group over the index group. Most units showed Gaussian-distributed firing rates about the mean firing rate for each movement, with 16 of the 60 units having at least one movement for which the normalized activations were not normally distributed (*p* < 0.001 for monkey N, *p* < 0.05 for monkey W, two-sided one-sample Kolmogorov-Smirnov test, corrected for false discovery rate).

**Table 1.** Summary of neural feature tunings to flexion versus extension.

| Monkey | Feature | Qty | Muscular Group |  |  |  | Untuned |
| --- | --- | --- | --- | --- | --- | --- | --- |
|  |  |  | Flexion | Extension | Opposed | Multiple |  |
| N | SU | 47 | 7 | 13 | 6 | 12 | 9 |
|  | SBP | 96 | 7 | 20 | 13 | 34 | 22 |
| W | SU | 13 | 0 | 0 | 0 | 8 | 5 |
|  | SBP | 96 | 2 | 23 | 5 | 51 | 15 |

The quantity (Qty) column indicates the total number of sources for each feature. SU means sorted units, SBP means spiking band power. Classifications were determined by $p < 0.001$ for monkey N or $p < 0.05$ for monkey W with a two-tailed two-sample Kolmogorov-Smirnov test between movement types. Opposed tunings showed activity levels different from baseline for opposite movements of the index and MRS fingers, where the two possibilities are index flexion with MRS extension or index extension with MRS flexion. Multiple tunings showed activity levels different from baseline for more than half but not all of the movements, indicating activation but perhaps not specificity. Decisions were made from 386 monkey N trials and 125 monkey W trials.

**Table 2.** Summary of neural feature tunings to index versus MRS group movements.

| Monkey | Feature | Qty | Finger Group |  |  |  | Untuned |
| --- | --- | --- | --- | --- | --- | --- | --- |
|  |  |  | Index | MRS | Both | Multiple |  |
| N | SU | 47 | 6 | 12 | 8 | 12 | 9 |
|  | SBP | 96 | 7 | 17 | 16 | 34 | 22 |
| W | SU | 13 | 0 | 0 | 0 | 8 | 5 |
|  | SBP | 96 | 10 | 3 | 17 | 51 | 15 |

The quantity (Qty) column indicates the total number of sources for each feature. SU means sorted units, SBP means spiking band power. Classifications were determined by $p < 0.001$ for monkey N or $p < 0.05$ for monkey W with a two-tailed two-sample Kolmogorov-Smirnov test between movement types. Here, multiple tunings showed activity levels different from baseline for more than half but not all of the movements, indicating activation but perhaps not specificity. Decisions were made from 386 monkey N trials and 125 monkey W trials.

In addition to performing all tuning analyses with the standard sorted units, we also included tuning analyses using the 300-1,000Hz spiking band power (SBP). We have previously shown that filtering spiking signals in the 300-1,000Hz band provides a signal that is highly correlated with the activity of the highest signal-to-noise ratio (SNR) units on an electrode (Nason et al., 2020). This is particularly effective for electrodes with low SNRs, as the 300-1,000Hz band was found to balance the tradeoff between signal and noise power (Irwin et al., 2016). Therefore, we constructed tuning curves from this band, which enabled us to perform the same analyses on sharper tuning curves from primarily multiunit electrode recordings. This resulted in more sources of unit activity than the limited quantities of sortable units on a Utah microelectrode array. As such, we nearly tripled the quantity of tuned neural features, from 26 of 60 tuned single units to 70 of 192 tuned SBP channels. Tuning curves for these channels are included in the left plots of each pair in Figure 5B, D, E, and F, with Table 1 and Table 2 showcasing similar preferences to sorted units. Most channels showed specificity to one muscle group and one finger group, with more representation of MRS movements than index movements in monkey N and vice versa in monkey W. Lastly, similar to the sorted units, all SBP channels showed Gaussian-distributed power levels about their respective means for all movements across all trials (*p* < 0.001 for monkey N, *p* < 0.05 for monkey W, two-sided one-sample Kolmogorov-Smirnov test, corrected for false discovery rate).

### Finger-Tuned Neural Activity Is Linear

To investigate how the neural activity relates to similar finger movements, we began with a classical cosine tuning analysis. Of the 41 tuned sorted units and 143 tuned SBP channels, 30 and 96, respectively, had significantly correlated sinusoidal fits (*p* < 0.05, Pearson’s correlation, corrected for false discovery rate). While sinusoidal tuning of a neuron to two-dimensional movements of a single limb may be reasonable (Georgopoulos et al., 1982), it may not be fundamental (Todorov, 2000), so the same rationale may not apply to movements of multiple limbs, or in our case, fingers. We have demonstrated that a linear Kalman filter enables online separation and individuation of two finger dimensions, so we sought to characterize the goodness of fit of several linear models.

To guide this process, we first calculated how well the activity of each movement could linearly predict the activity of each other movement. Figure 4 illustrates this, with more yellow (or blue) cells indicating greater (or more negative) coefficients for the activity on the horizontal axis to predict the activity on the vertical axis. Generally, we found that the activity corresponding to most movements could be strongly predicted by the activity of its most related movements (the yellow off-diagonal bands). For example, the activity corresponding to MRS-only flexion (MF) can be predicted significantly by the activities corresponding to MRS flexion/index extension (MI) and index & MRS flexion (F). This suggests two potential linear models. The first makes the naïve assumption that the neural activity of a given movement can be predicted by averaging the activity of its component movements (Avg), and the second generalizes the first by computing a weighted average of the activity of the component movements with coefficients fit via regression (LR). Additionally, the off-diagonal green bands in Figure 4 that represent lower, but non-zero coefficients suggest some predictable information may be contained in the activity of opposite movements (i.e. index flexion is predictive of index extension). Consequently, the third linear model expands LR to include the activity of the opposite movement, also fit via regression (LRO). The left plots of each pair in Figure 5 show the raw data with an overlaid tuning curve, and the right plots of each pair overlay the activity predictions from each model onto the true activity. Visually, all models predicted most activity well, with most errors occurring at the peaks/tuning preferences of a given unit or SBP channel (SBP included due to its specificity to single units; Nason et al., 2020). Figure 6 illustrates the fit statistics for the different models. First, in Figure 6A, the Avg model achieved the lowest number of predicted tuning curves significantly correlated with the actual tuning curves, with just 13 of the 181 total sorted units and SBP channels (*p* < 0.05, Pearson’s correlation, corrected for false discovery rate). Alternatively, the LR and LRO models provide a better fit, predicting 105 and 113 significantly correlated tuning curves of the total 181, respectively. Between LR and LRO, information about the opposite movement did not substantially improve the quantity of units and SBP channels whose tuning curves could be significantly predicted. Thus, the drastic improvement of the LR and LRO models over the Avg model suggests that the finger-related neural activity for any particular movement may not be equally representative of the component movements despite reliably strong activation for those component movements. Looking at the plots of predicted versus actual activity in Figure 6C, we see more evidence of the same result: lower errors that are normally distributed about zero (*p* < 1 × 10^−10^, two-sided one-sample Kolmogorov-Smirnov test) and tighter binding to a unity line for the regression models. The weights for the regression models are summarized in Supplementary Figure 2.

**Figure 4.**
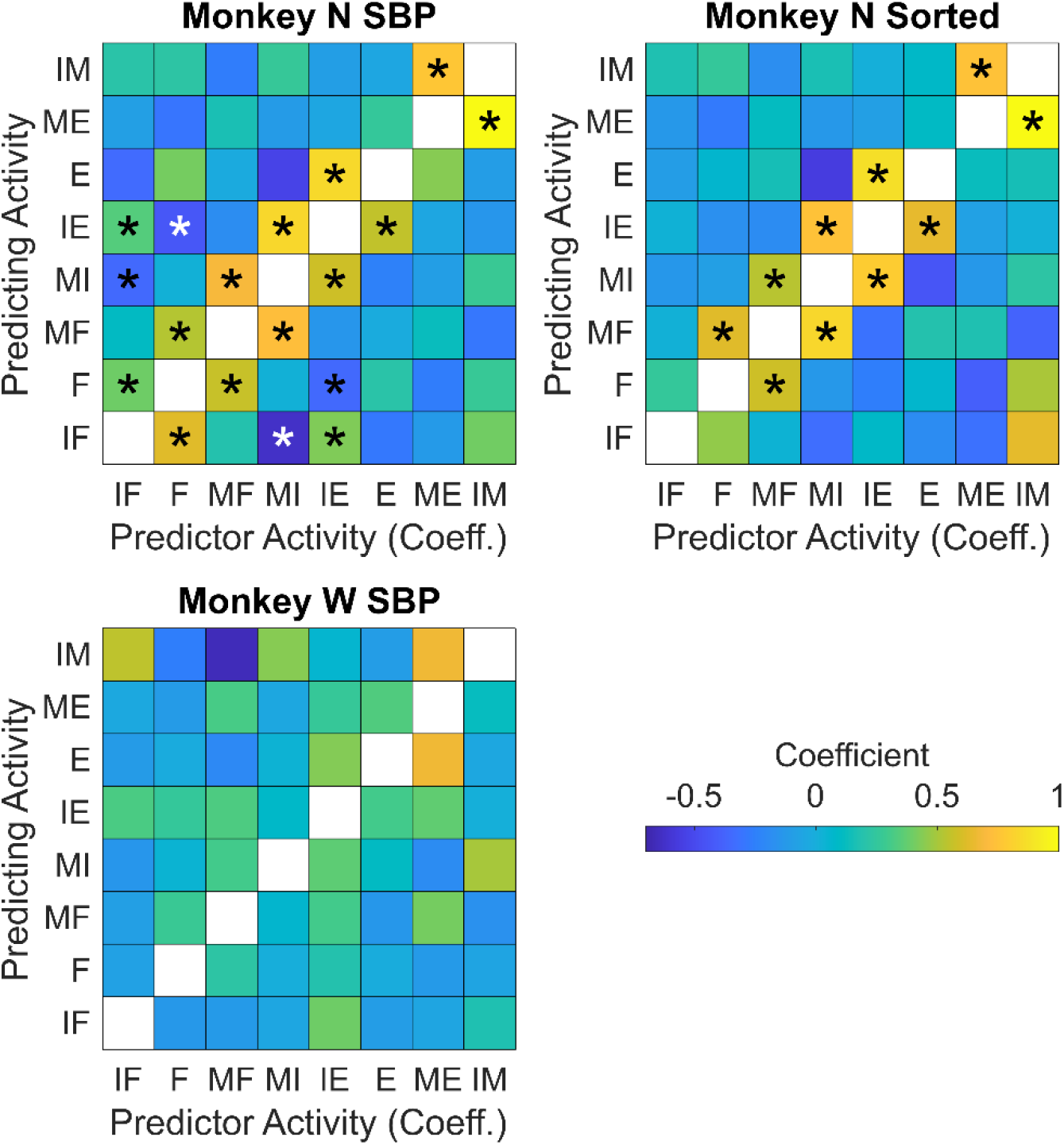
Predictability of the activity of each movement from the activities of other movements. The top and bottom rows of axes represent monkeys N and W, respectively, and the left and right columns of axes represent SBP and sorted unit results, respectively. A cell in each set of axes is colored based on the predictor’s coefficient (horizontal axis) when predicting the activity of a movement (vertical axis). Asterisks indicate statistical significance (based on each coefficient’s t-score, *p* < 0.001).

**Figure 5.**
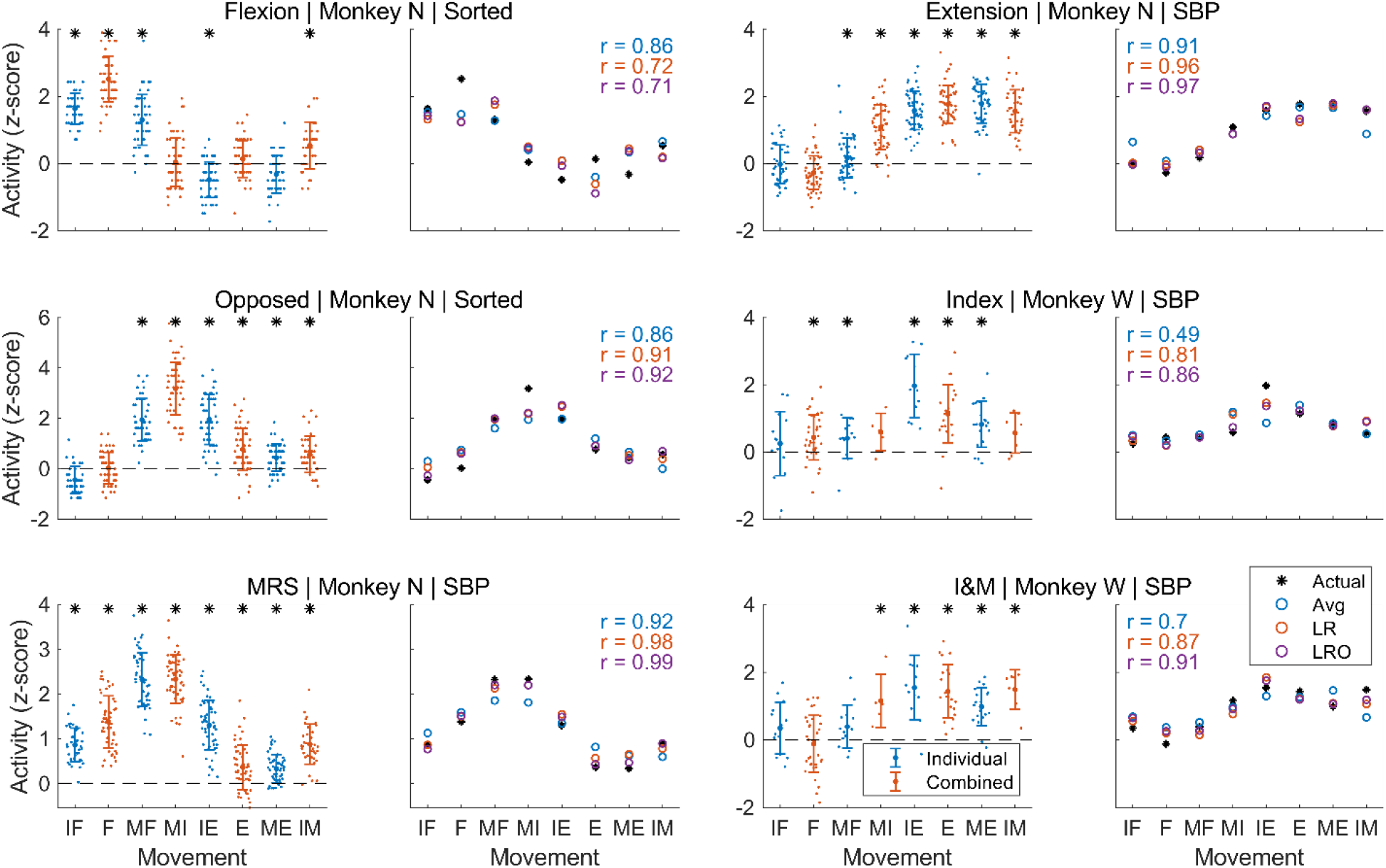
Exemplary tuning curves and linear predictability of the activity. The text above each pair of plots first indicates the tuning preference of a sorted unit or SBP channel, followed by the subject from which the activity was recorded, then the feature type. I&M suggests preference to both the index and MRS groups. The left plot of each pair displays the true tuning curve of the given sorted unit or SBP channel. The large dot and error bars represent the mean and standard deviation for the peri-movement activities across all trials of each type of movement. The smaller dots represent the peri-movement activity for each trial of each type of movement. Asterisks indicate significant difference from the unit’s or channel’s mean activity, or zero z-score, across the experiment (two-sided two-sample Kolmogorov-Smirnov test, *p* < 0.001 for monkey N, *p* < 0.05 for monkey W, corrected for false discovery rate). Individual group movements are plotted in blue, combined group movements are plotted in orange. The right plot of each pair displays the mean activity for each movement (copied from the left plot in asterisks) as well as the predictions of the activity for each movement. For example, for IF in C, the black asterisk is the mean dot copied from the left hand plot, the blue circle (Avg) is the average of the asterisks for F and IM, the orange circle (LR) is the weighted sum of the asterisks for F and IM, and the purple circle (LRO) is the weighted sums of the asterisks for F, IM, and IE. Pearson’s correlation coefficients between the predictions and the true activity are displayed in each plot.

**Figure 6.**
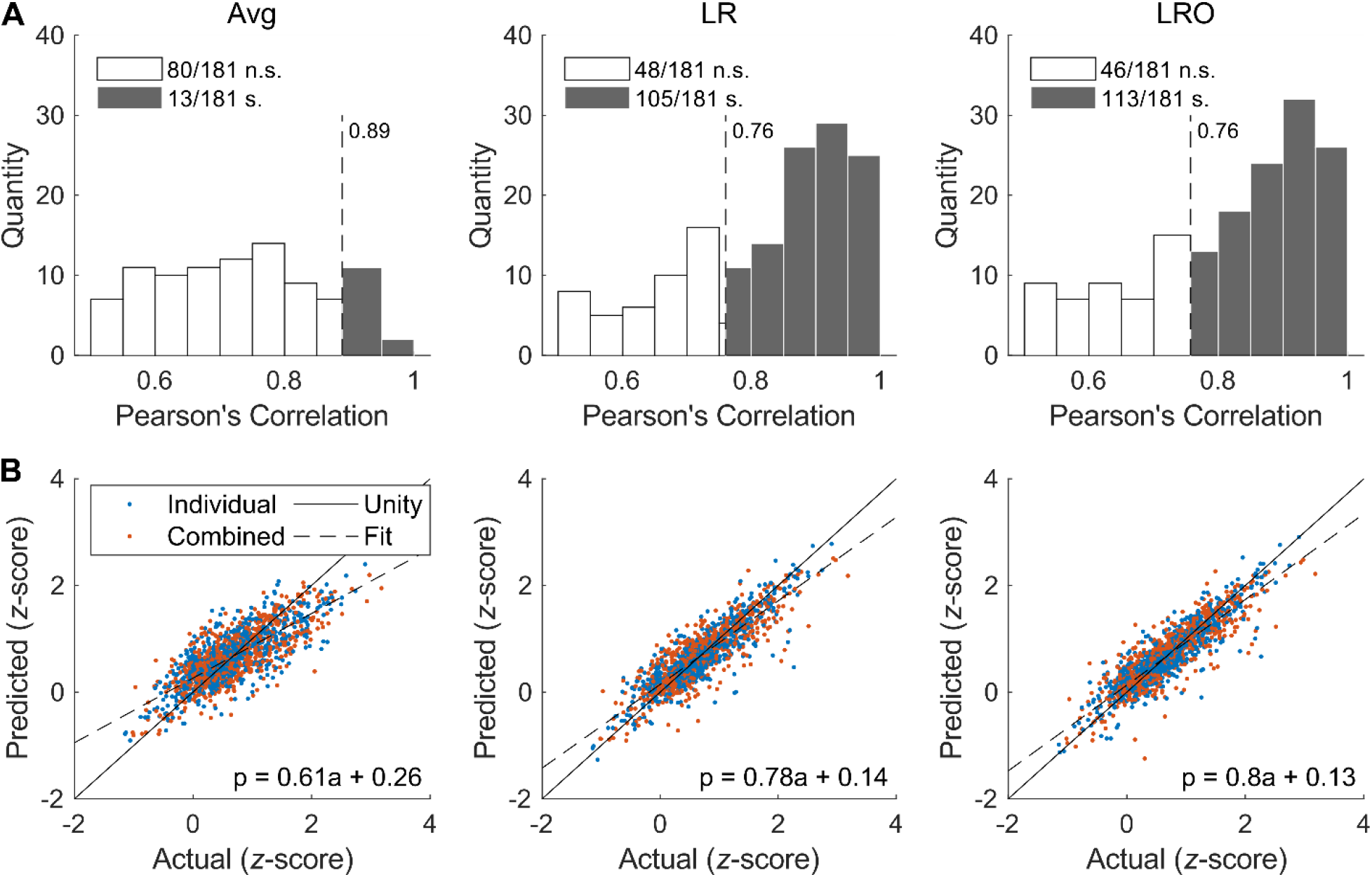
Statistics of the linearity in tuning curves. Sorted units and SBP channels from monkey N and SBP channels from monkey W that had at least one movement significantly different from the average activity and an average firing rate greater than 2Hz (for sorted units) are included in analysis. The first column represents results from averaging the neighbors, and the second and third columns represent results from linearly regressing the neighbors without and with the opposite movement, respectively. (**A**) Histogram of the correlation coefficients (Pearson’s r) between the predicted tuning curves and the actual tuning curves. The histogram is limited to the 0.5-1 range as we found this range of tuning curves to be visually linear. The dotted line with the attached number indicates the cutoff for significance after correction for false discovery rate. n.s. means not significant, s. means significant with *p* < 0.05. (**B**) Scatter plots of the predicted activity vs. the true activity for each movement. Blue dots represent individual group movements and orange dots represent combined group movements. The solid line is unity, where the predicted activity would equal the true activity. The dashed line is fit to the data.

### Linear Models Generalize to Predict Untrained Finger Movements

From a decoding perspective, it is helpful to know the training data requirements for a high performance, multidimensional hand neural prosthesis. Specifically for our two-finger task, in addition to addressing the linearity of neural activity, might a decoder model need to be trained on both individual and combined finger group movements, or is the neural activity for one of the types of movements sufficient to generate a model that encompasses both?

To answer that question, we split the trials (1,018 from monkey N and 368 from monkey W) into two subsets: one representing individual finger group movements only and the other only combined movements of both finger groups. Then, we trained linear regressions with ridge regularization on each set of trials to predict finger positions using exclusively sorted units or SBP and tested them on the corresponding sorted units or SBP of the untrained set of trials. For example, we first trained regressions on *individual* movements of the index or MRS finger groups and used the trained algorithm to decode the *combined* movements of the index and MRS finger groups, and vice versa. Figure 7 illustrates the averaged predicted traces from split training for all trials of a given movement overlaid on the averaged predicted traces for that same movement using a decoder trained on all trials with cross-validation. Monkey N’s traces show that the split-trained decodes mostly overlay the full-trained decodes, which suggests that finger individuation information in the population’s neural activity is preserved across movements, even for untrained behaviors.

**Figure 7.**
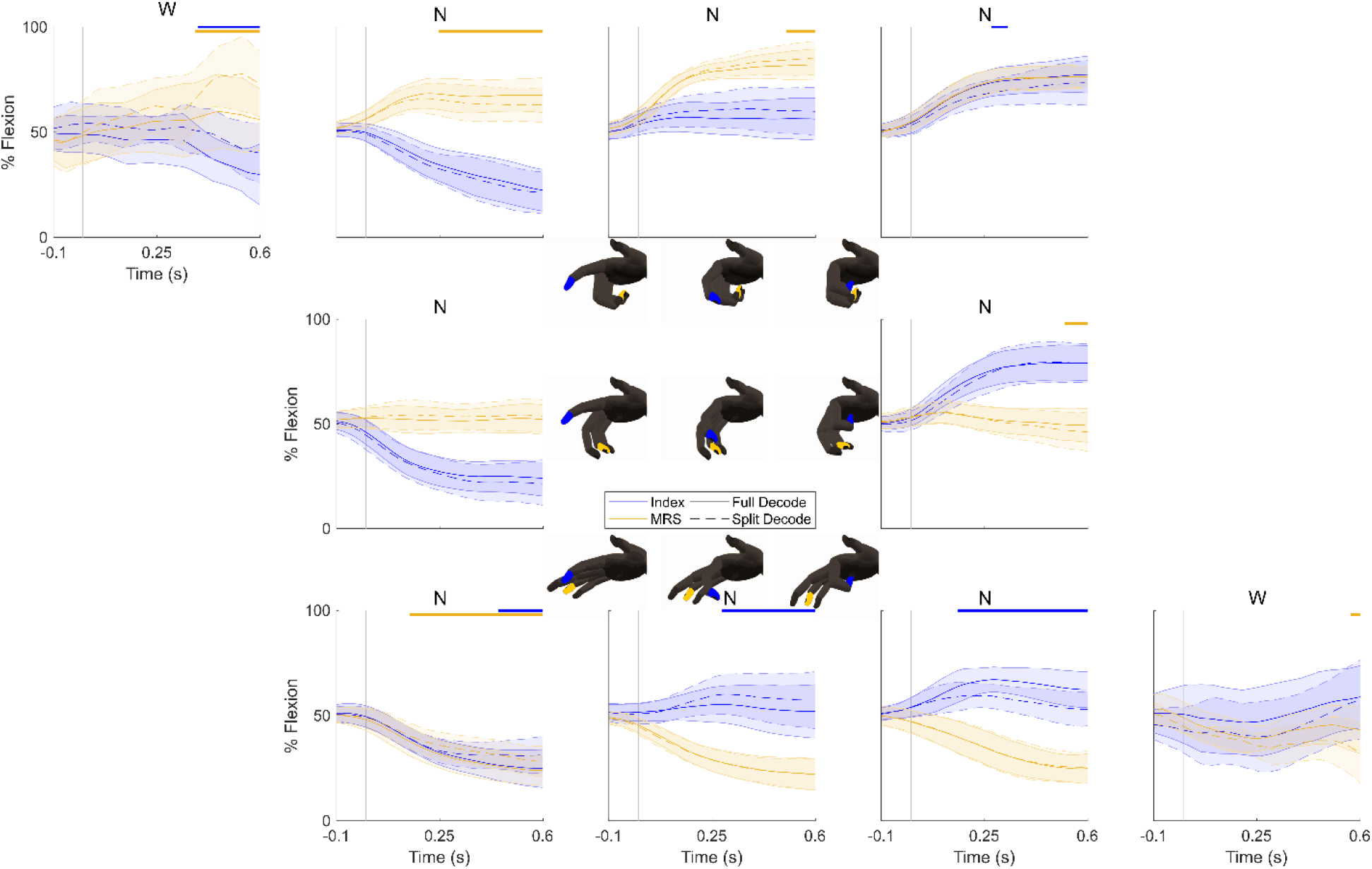
Offline ridge regression decoding of all SBP channels, trained on either individual or combined finger group movements. The central trace is the average predicted behavior from all trials of the indicated movement and the shaded region is the standard deviation, all aligned in time by the movement onset (vertical gray line). Individual finger group movements (four middle plots) were decoded using a regression model trained on combined finger group movements (four corner plots), and vice versa for the combined finger group movements. These are represented by the “Split Decode” dashed traces. The “Full Decode” solid line traces represent the average decode given the full dataset to train the regression model, with cross-validation. The blue traces correspond to the index group and the yellow traces correspond to the MRS group. The yellow or blue lines near the top of each plot indicate significant differences between the two mean predicted positions based on a bootstrap analysis on the differences (greater than a one-sided 95% confidence interval). The cartoon hands near each plot demonstrate the movement being decoded, and the letter above each plot indicates which monkey’s decodes are displayed. The plots in the top left and bottom right corners are the corresponding decodes from monkey W, included to show that individuation of the finger groups is possible with a decoder model trained on individual finger group movements despite low signal-to-noise ratio neural activity.

However, there are differences during some of the predictions (greater than a one-sided 95% confidence interval of a bootstrap analysis on the errors). Differences in these predicted traces align with the results found during the tuning curve analyses, where the neural activity of particular movements requires unequal contributions of the component movements. As such, by the assumption of linearity, there should be some loss in performance associated with testing on untrained behaviors, which is shown in Figure 7.

Although monkey W had limited representation of MRS movements, the better-represented index finger neurons enabled the prediction of opposed finger movements (IM and MI), despite the regression not being trained on those behaviors. Thus, they demonstrate the same result, that individuation in the spiking activity is preserved for untrained behavior. We included the full set of averaged decodes for the eight movements using sorted units and SBP for both monkeys in Supplementary Figures 3-6. To avoid any confounds from averaged traces, we also included plots of the individual trials’ decodes used to calculate the averages in Supplementary Figures 7-10. We also performed the same analysis for each unit and SBP channel, which is summarized in Supplementary Figure 11.

## DISCUSSION

Modern hand neural prostheses have not yet been able to reproduce individuated finger movements across their entire ranges of motion. Here, we demonstrated that a linear Kalman filter with or without intention retraining is adept at decoding individuated movements of an index finger and a middle-ring-small finger group in real-time with high-performance, given sufficient number of tuned units. We then showed that monkey N could improve his control performance using the ReFIT Kalman filter. To address how this complex task can be explained with a linear decoder, we found that the neural activity of any particular movement can be reliably predicted from the activity of its related movements via a weighted sum, providing an explanation for our high performance linear decoding model. Finally, we validated this claim by showing that combined finger group movements can be predicted with a decoder model trained on individual finger group movements and vice versa. Only a slight performance loss was realized when comparing predictions of decoders trained on the full set of behaviors and decoders trained only on trials representing individual or combined finger movements.

The complexity and novelty of our 2D task proposes interesting questions regarding the interpretations and intentions of the monkeys performing it. We proposed two frameworks for retraining linear Kalman filters based on the monkeys’ intentions, one assuming each finger is its own task for two total objectives and the other assuming both fingers belong to one two-dimensional task. While both improved decoding performance substantially, neither outperformed the other despite making clearly different assumptions about the monkeys’ interpretations of the task. As such, we think these data suggest two possible truths: the nuances of intention-based recalibration methods for the prediction of continuous two-finger movements will not significantly impact performance, or the optimal recalibration scheme for multiple finger movements remains unknown.

By applying the definition of linearity to our experiments in which we trained decoders on individual finger movements and used them to predict combined finger movements, we did not expect the positions predicted by the split decoder and the positions predicted by the full decoder to be so similar, which they were in many cases. We propose a few potential explanations. First, this could be attributed to using behaviors of different magnitudes for each direction, such that when plotting just the middle magnitude, the differences in predictions were negligible. Second, since the tuning curves showed graded activations to several related movements, combinations of the contributions from different units and SBP channels may be sufficient to predict movements of greater magnitude. Third, finger-related neural activity may have a minor non-linearity, either between units and SBP channels or with its relationship to the behavior, which has been reported previously (Naufel et al., 2019).

Our real-time decoding results suggest that linear models can accurately fit the movements of two independent fingers across their full ranges of motion during online control. Although our initial individual finger datasets suggested that this may require nonlinear models (Vaskov et al., 2018), the results presented in this manuscript indicate otherwise for combined movements. Sufficiency of linear models has major implications for clinical neural prostheses. It has been previously suggested by unconstrained finger movements that primary motor cortex linearly predicts movements of the hand and fingers in monkeys (Aggarwal et al., 2013; Ethier et al., 2012; Kirsch et al., 2014; Okorokova et al., 2020) and humans (Ajiboye et al., 2017; Bouton et al., 2016; Wodlinger et al., 2015) when many of the DoFs may move along very similar trajectories. This work extends those results to two well-separated independent finger DoFs and suggests that this will be possible in current human clinical trials studies. Finger-related neural activity in primary motor cortex recorded with human-grade Utah microelectrode arrays can be sufficient to individuate at least two systematically separate DoFs using linear models. Further, linear individuation of two DoFs within the hand, as shown here, suggests the possibility of linear models sufficiently individuating more of the 27 DoFs within the hand, as has been hinted for low-magnitude movements (Kirsch et al., 2014).

In terms of clinical viability of a neural prosthesis, where usage outside of a laboratory, hospital, or rehabilitation environment is the ultimate goal, the computational simplicity and generality of linear models make them promising solutions. Decoders such as the Kalman filter, Wiener filter, or ridge regression (Collinger et al., 2013; Ethier et al., 2012; Malik et al., 2011) require surprisingly few computations per iteration, opening the possibility of implementation on portable or implantable devices. Additionally, our results suggesting that the activities of individual movements can be linearly combined into the activities of more complex movements hint that decoders may not need to be trained on the full suite of behaviors that they will be used to predict. Instead, training decoders on orthogonal behaviors that span the full behavioral space, with representative neural activity, may be all that is required. This may cut the 5-10 minutes of decoder training time drastically, potentially streamlining the daily calibration of an outside-of-laboratory neural prosthesis. Generally, the results presented here suggest that naturalistic hand and finger neural prostheses with many DoFs may be close to clinical translation using simple linear decoder models and simple training procedures.

Though the results presented here show that linear decoding models can predict the movements of individuated fingers with high performance, the decoders were unable to achieve the level of precision and control of the able-bodied hand. To bridge that gap, decoders may need to account for a nonlinear relationship between cortical activity and behavior. Several nonlinear neural networks have been tested for brain-machine interfaces (Hosman et al., 2019; Pandarinath et al., 2018), though few have transitioned to testing online. An early online recurrent neural network (Sussillo et al., 2012) showed that neural architectures have promise in online decoding. However, due to the heavy computational requirements of neural networks, they must be optimized and significantly compressed before being considered for out-of-laboratory, portable, and implantable brain-machine interfaces. Therefore, it is valuable to characterize the limits of linear models in discriminating neural states with truly simultaneous movement of independent DoFs, as was presented with fingers in this work.

Importantly, the linearity of finger-related neural tuning models gives insight into how primary motor cortex represents the activity of a variety of related movements. Most neural units presented in this study did not show nonlinear specificity to one movement, but graded tuning to several related movements. In fact, most of the tuning curves fit the classical cosine tuning model demonstrated for arm reaches (Georgopoulos et al., 1982) and finger movements (Georgopoulos et al., 1999). Despite directional tuning with arm reaches fitting logically (the angles and magnitudes of movement can all be referenced to one limb in a radial task), the physiological assumptions are broken by our task with two independent finger dimensions that have their own relatively independent muscles. Our task employs what are essentially two limbs (or fingers) traversing their own spaces with any given neuron capable of being tuned to both limbs, making musculoskeletal models seem like the more relevant explanation for the underlying neural activity (Todorov, 2000). While it is intuitive to conclude that the cosine tuning model expands to our two-finger task, we think our weighted-average model better explains how neurons can simultaneously encode movements of multiple independent limbs.

This raises a major question: how far can this weighted-average model be extended as we consider more DoFs? Several groups have investigated how neurons tuned to multiple behaviors respond to tasks requiring those behaviors (Cross et al., 2020; Diedrichsen et al., 2013; Heming et al., 2019; Jorge et al., 2020; Stavisky et al., 2019, 2020; Willett et al., 2020). One way to explain this is that the neural activity underlying separate and combined behaviors can be explained by multiple orthogonal subspaces. Here, we have evidence to suggest that linear models can effectively combine at least two finger subspaces that are substantially related. Provided sufficient quantities of neurons representing those subspaces, we think compound movements may continue to be well represented by linear combinations of their component subspaces, up to the full dimensionality of the hand and beyond to the entire motor system.

## METHODS

All procedures were approved by the University of Michigan Institutional Animal Care and Use Committee.

### Array Implants

We implanted two male rhesus macaques, age 7 at the time of data collection, with Utah microelectrode arrays (Blackrock Microsystems, Salt Lake City, UT, USA) in the hand area of primary motor cortex, as described previously (Irwin et al., 2017; Vaskov et al., 2018). Pictures of the implants are illustrated in Figure 1D. Only motor cortex arrays were used in this study.

### Feature Extraction

All processing was done in Matlab versions 2012b or 2018a (Mathworks, Natick, MA, USA), except where noted. Threshold crossing rates were processed and synchronized in real-time during the experiments (see the subsequent section for a description of data flow). We configured the Cerebus neural signal processor (Blackrock Microsystems) to extract voltage snippets that crossed a −4.5 times the root-mean-square (RMS) threshold, customized to each channel. Then, these waveforms were streamed to a computer running xPC Target version 2012b (Mathworks), which logged the source channel of each spike and the time of each spike’s arrival relative to all other real-time experimental information. Both monkeys had 96 channels of threshold crossing rate data analyzed, though for closed-loop decoding, channels were masked to those that were not clearly disconnected and had contained morphological spikes during the experiment or at some time in the past (see SBP section below for reasoning).

We also extracted sorted unit firing rates for offline analyses. We imported the relevant broadband (0.1Hz – 7.5kHz sampled at 30kSps) recordings into Offline Sorter version 3.3.5 (Plexon, Dallas, TX, USA). Then, we high-pass filtered the recordings with a 4-pole Butterworth filter with a cutoff frequency set to 250Hz. To sort clear units, we used the threshold level determined during the experiment by the Cerebus at −4.5RMS, then eliminated clearly artifactual threshold crossings. Then, we sorted the remaining spikes crossing the threshold individually and in combination with principal component analysis, clusters as determined by k-means or Gaussian mixture model clustering (as implemented in Offline Sorter), and visual inspection. The spike timings of each sorted unit were then re-synchronized with the experimental data offline. After all sorting, monkey N had 47 units and monkey W had 13 sorted units.

Spiking band power was also acquired in real-time by the same experimental system as threshold crossing rates. We configured the Cerebus to band-pass filter the raw signals to 300-1,000Hz using the Digital Filter Editor feature included in the Central Software Suite version 6.5.4 (Blackrock Microsystems), then sampled at 2kSps for SBP. The continuous data was streamed to the computer running xPC Target, which took the magnitude of the incoming data, summed all magnitudes acquired in each 1ms iteration, and stored the 1ms sums as well as the quantity of samples received each 1ms synchronized with all other real-time experimental information. This allowed offline and online binning of the neural activity with 1ms precision. As with threshold crossing rate, we masked channels for closed-loop decoding to those that were not clearly disconnected and had contained morphological spikes during the experiment or at some time in the past, as SBP could possibly extract firing rates of low-SNR units remaining represented on such channels (Nason et al., 2020).

### Experimental Setup

The experimental apparatus used for these experiments is the same as described previously (Irwin et al., 2017; Nason et al., 2020; Vaskov et al., 2018). Briefly, the monkeys’ Utah arrays were connected to the patient cable (Blackrock Microsystems) and raw 0.1Hz-7.5kHz unfiltered broadband activity at 30kSps, 300-1,000Hz activity at 2kSps, and threshold crossings at a −4.5RMS threshold were extracted from the neural recordings by the Cerebus for storage. The 2kSps and threshold crossing features were streamed to the xPC Target computer in real-time via a User Datagram Protocol packet structure. The xPC Target computer coordinated several components of the experiments. It binned threshold crossings and SBP in customizable bin sizes, coordinated target presentation, acquired measured finger group positions from one flex sensor per group (FS-L-0073-103-ST, Spectra Symbol, Salt Lake City, UT, USA), and transmitted finger positions along with target locations to an additional computer simulating movements of a virtual monkey hand (MusculoSkeletal Modeling Software) (Davoodi et al., 2007). Task parameters, states, and neural features were stored in real-time for later offline analysis.

### Behavioral Task

We trained monkeys N and W to acquire virtual targets with virtual fingers by moving their physical fingers in a more complex version of the two-finger task we published previously (Nason et al., 2020). During all sessions, the monkeys sat in a shielded chamber with their arms fixed at their sides flexed at 90 degrees at the elbow, resting on a table. Monkey N had his left hand and monkey W had his right hand placed in the manipulandum described previously (Vaskov et al., 2018). The monkey sat in front of a computer monitor displaying the virtual hand model and targets described previously. The monkeys were trained to move their MRS fingers as one group independently from the index finger as its own group, and the fingers of the virtual hand were actuated accordingly.

Each trial began with one spherical target appearing along the one-dimensional movement arc of each finger group, for a total of two simultaneous targets. Each target occupied 15% of the full arc of motion of the virtual fingers. Targets were presented in a classical center-out-and-back pattern when viewed from the two-dimensional behavioral space illustrated in Figure 1B (Georgopoulos et al., 1982). Every other target was presented at a rest position, 50% between full flexion and full extension. The non-rest targets had a direction (0, 45, 90, 135, 180, 225, 270, or 315 degrees, corresponding to index flexion, both flexion, MRS flexion, MRS flexion with index extension, index extension, both extension, MRS extension, and index flexion with MRS extension, respectively) and magnitude (±20%, ±30%, or ±40% flexion from rest, except for 135 and 315 degrees, as ±40% was too far of a split in the finger groups to be performed reliably) pseudo-randomly chosen in the two-dimensional finger space to generate the two targets in a center-out-and-back pattern. The presentation order of the targets was random, though the same order was repeated across sets of trials. Approximately one year after data collection, several experiments with a true-random order of the targets validated no change in performance. For a successful trial, the monkey was required to move the virtual fingers into their respective targets and remain there for 750ms continuously in manipulandum control or 500ms continuously in brain control (see below). For training and testing of closed-loop decoders, 135 and 315 degrees posture styles were not included as target presentation options. Upon successful trial completion, the monkeys received a juice reward.

### Neural Activity Normalization

All of our tuning curve analyses used normalized neural activity. For each sorted unit or SBP channel, we computed the mean, standard deviation, and root-mean-square activity level across the entire experiment in 300ms bins. Then, we eliminated all trials that were not center-to-out, that were unsuccessful, or had a duration longer than 3.0s, indicating possible distraction, struggle, or other factors that may confound tuning analyses.

For each trial, we estimated movement onset by the following procedure. First, we smoothed the positions, measured at 1kHz but down-sampled 20-fold, with a 2^nd^ order, 11 frame-length Savitzky-Golay filter using Matlab’s *sgolayfilt* function, as was done previously (Nason et al., 2020). The movement onset for each trial was then calculated as the first negative peak in speed just before the first positive peak in speed for any finger group that moved to a non-rest target. We found this procedure matched estimated movement onsets visually well for nearly all observed trials.

Given the movement onset, then for each included trial, we extracted the activity for each sorted unit or SBP channel from 100ms prior to 200ms after movement onset. We then normalized these measurements by subtracting th e unit’s or SBP channel’s mean then dividing by the standard deviation. This results in one normalized level of activity for each unit or SBP channel for each trial.

### Computation of True and Predicted Tuning Curves

We calculated tuning curves for each monkey using one isolated experiment during which monkey N performed 2,130 and monkey W performed 969 manipulandum control trials of the center-out-and-back task. After eliminating trials that were not center-to-out, that were unsuccessful, had a duration longer than 3.0s, or had computed movement onset times (see prior section) less than 100ms into the trial, monkey N had 1,018 and monkey W had 364 remaining trials. All of these trials were used for the ridge regression analyses (see section below), while only trials in which the monkeys moved their fingers ±30% from rest were used for tuning analyses.

To compute sinusoidal tuning curves, we fit the amplitude, period, phase, and offset by using Matlab 2018a’s fminsearch function minimizing the total squared error over a maximum 1,000,000 iterations.

We tested three models of predicting the firing rates of all units corresponding to some movement given the activity of the units for other movements. The first naïvely assumes that the firing rates corresponding to one movement is the average of the firing rates of its component movements. The second and third assume that the firing rate *n* of some movement *m* is a weighted sum of the firing rates corresponding to the other movements along with some constant offset *C*_0_. This can be modeled by the following linear equation for one channel with an arbitrary *k* number of movements:

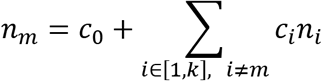

Let *n*_*m,u*_ be the firing rate from neural unit *u* for movement *m*. Then *N*_*m*_ is the vector of firing rates from all neural units corresponding to movement *m* and *N*_*D*_ is the training data matrix containing firing rates from all neural units corresponding to all of the movements that are not movement *m*:

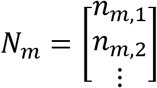

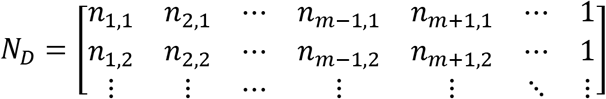

As such, we can solve for the vector of coefficients for one movement *m* by solving the following linear regression equation using the data given by all valid units:

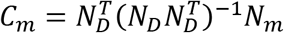

The first of the two regression models we tested assumed that *n*_*m*_ is composed of some weighted amounts of its component movements, which are *n*_*m*−1_ and *n*_*m*+1_ in this formulation, with a learned offset. The second model assumed that *n*_*m*_ is composed of some weighted amounts of *n*_*m*−1_, *n*_*m*+1_, the learned offset, and the firing rates corresponding to the opposite movement. We performed these same procedures using SBP in place of the firing rates.

We predicted the neural activity for each movement based on the two regression models with leave-one-out cross-validation on the neural sources. The cross-validated linear models for predicting sorted unit activity were trained only on the other sorted units within each monkey. The same was done for SBP channels. Sorted units and SBP channels that had no significantly tuned movements according to a two-sample two-tailed Kolmogorov-Smirnov test (*p* > 0.001 for monkey N, *p* > 0.05 for monkey W) and sorted units with mean firing rates under 2Hz were excluded from analyses. Monkey W’s sorted units were also excluded from some analyses, as only three of the units showed any significant tuning, forcing us to rely more on SBP for this monkey. This left a total of 133 SBP channels (84 from monkey N and 49 from monkey W) and 38 sorted units (from monkey N).

### Decoding of Neural Activity

We executed two different algorithms to predict finger movements from neural activity.

#### Closed-Loop Kalman Filtering

First, to assess neural prosthetic performance in application, we gave the monkeys visual feedback of the decoders’ outputs during the behavioral task. For each closed-loop experimental session, the monkeys began by completing at least 350 trials with the virtual hand controlled directly by the movements of the manipulandum. The monkeys were required to acquire and hold the targets for 750ms continuously for a successful trial with a 10s trial timeout. The behavioral data (i.e. one-dimensional positions per finger group) were measured synchronously with the non-normalized neural features by the xPC Target computer. Then, we trained a standard position/velocity Kalman filter on this data binned at 32ms (which we found superior to other tested bin sizes), as described previously (Irwin et al., 2017), using Matlab version 2012b (Mathworks). For predictions of two finger dimensions, the Kalman filter assumed a kinematic state of one position and one velocity for each group:

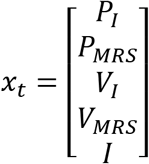

The Kalman filter predicts the state at each timestep based on an optimal combination of two different predictions. The first is a prediction made based on the state of the previous timestep, and the second is a prediction made based on a comparison between the measured neural activity and that predicted by the predicted kinematics of the current timestep. This can be summarized by the following equations:

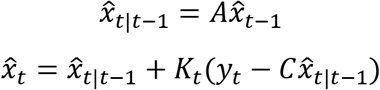

where *y*_*t*_ is a vector of neural features at the current time step, *K*_*t*_ is the Kalman gain balancing how much the neural activity should contribute to the final prediction, *A* is the state transition matrix, and *C* is a linear regression trained to convert kinematics to neural features. Training of these matrices was performed as described previously (Irwin et al., 2017) but extended to account for both index and MRS. The *A* matrix was fit to take the following form:

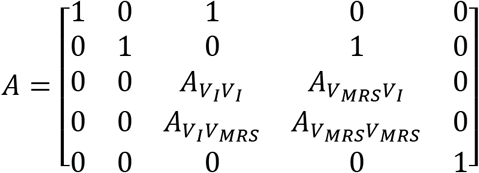

After training the standard Kalman filter, we computed the Kalman filter’s predictions in real-time to actuate the virtual hand independent of the monkeys’ physical movements. For successful acquisition, the monkeys were required to acquire and hold the targets for 500ms continuously with a 10s trial timeout for monkey N or a 30s trial timeout for monkey W. The positions of the virtual fingers were computed by integrating predicted velocities over time, with an initial position taken as the true position of the fingers before entering brain control mode. Monkey W only performed closed-loop experiments using the standard Kalman filter, while monkey N used both the standard Kalman filter and the dual-stage training ReFIT Kalman filter. To train the ReFIT Kalman filter, monkey N used the standard Kalman filter in closed-loop for at least 250 trials. Then, we used one of three intention-training paradigms to train the ReFIT Kalman filter.

1. The first is a two-dimensional form of the ReFIT Kalman filter we previously proposed (Vaskov et al., 2018), where we negated the incorrectly predicted one-dimensional velocity for each finger and set the velocities of fingers within their targets to zero before recomputing the regression matrices. This option assumes the monkey viewed the task as two separate one-dimensional problems and is highly related to the visual cues.
2. The second training method is very similar to the original ReFIT Kalman filter training method (Gilja et al., 2012). In the two-dimensional behavioral space illustrated in Figure 1B, we rotated the net two-dimensional velocity towards the target’s center for every trial and calculated the component index and MRS velocities from the net velocity. This option assumes the monkey viewed the task as one two-dimensional problem and is highly related to our two-dimensional space in which the two one-dimensional targets are mapped to one two-dimensional target.
3. The third training method combines the first two. We concatenated the recalibrated predictions from each method to create a 2*n* × *d* predictions matrix, where *n* is the number of timesteps in the training data and *d* is the dimensionality of the behavior (5 in this case). Then, we concatenated one copy of the *n* × *f* neural feature matrix (*f* being the number of features) to match the size of the first dimension of the predictions matrix to recompute the regressions. This option assumes the monkey viewed the task as some combination of two one-dimensional tasks and one-two dimensional task, which reflects some of the behavior seen in Supplementary Figure 12.

After training the ReFIT Kalman filter, we again computed the predictions in real-time using the new model and delivered visual feedback of the predictions to the monkey by actuating the virtual hand.

#### Open-Loop Ridge Regression

When trying to gauge the generalization of linear models to untrained finger behaviors, we used ridge regression (with a regularization term *λ* = 0.001) as it does not depend on iteration for continued stability. The neural activity was binned every 100ms and 10 total bins were used per feature: one for the current time step and one for each of the previous 9 timesteps for a total 1s of neural data. Neural activity from before each trial’s beginning was assumed to be 0 for sorted units and each channel’s root-mean-square value for SBP, as had been done previously (Chestek et al., 2007).

After sorting all trials as previously described, we split them into two sets: individual and combined finger group movements. Then, we trained a regression model on one set and tested on the other. Additionally, we performed 10-fold cross-validation on the full dataset to gauge information loss without training on the full set of behaviors. To compare these, we performed a bootstrap analysis on the errors between the two decodes for all time points in each experiment. Then, if the error for any averaged sample was greater than the upper one-sided 95% confidence interval resulting from the bootstrap analysis, the split decode was deemed significantly different from the full decode.

#### Performance Metrics

To evaluate performance, we used two metrics. First, target acquisition time was computed as the total time between a trial’s beginning and its conclusion after successfully holding the target for the indicated period, less the hold time. We opted to exclude the hold time from the calculation so that the computed metrics better represented the time taken to reach the targets. Note that we required the monkey to hold targets for longer time periods when in manipulandum control mode so that there would be sufficient representation of neural data corresponding to stopping when training the online decoder.

The second performance metric is throughput, measured in bits per second by way of Fitts’ Law (Fitts et al., 1954). Fitts’ Law has had many interpretations beyond its original one-dimensional form since it was originally introduced. In the case of our behavioral task, the two dimensions each have their own one-dimensional target. As such, we found the following interpretation of Fitts’ Law to best represent the simultaneous completion of two one-dimensional tasks:

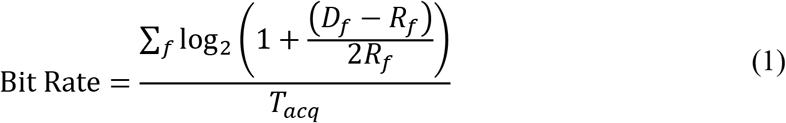

where *D*_*f*_ is the distance between the starting point of finger *f* and the center of its target, *R*_*f*_ is the radius of the target for finger *f*, and *T*_*acq*_ is the time taken to acquire the target less the hold time. Such a formulation gives an independent index of difficulty for each target, as was proposed by others investigating multiple target tasks (Stoelen and Akin, 2010). In our behavior task, for any finger group whose starting position is already within its target (i.e. in the case of a center-to-out movement of just one finger group), the index of difficulty for that finger is 0.

## ACKNOWLEDGEMENTS

We thank Eric Kennedy for animal and experimental support. We thank Gail Rising, Amber Yanovich, Lisa Burlingame, Patrick Lester, Veronica Dunivant, Laura Durham, Taryn Hetrick, Helen Noack, Deanna Renner, Michael Bradley, Goldia Chan, Kelsey Cornelius, Courtney Hunter, Lauren Krueger, Russell Nichols, Brooke Pallas, Catherine Si, Anna Skorupski, Jessica Xu, and Jibing Yang for expert surgical assistance and veterinary care. This work was supported by NSF grant 1926576, Craig H. Neilsen Foundation project 315108, A. Alfred Taubman Medical Research Institute, NIH grant R01GM111293, and MCubed project 1482. S.R.N. was supported by NIH grant F31HD098804. M.J.M. was supported by NSF grant 1926576. A.K.V. was supported by fellowship from the Robotics Graduate Program at University of Michigan. M.S.W. was supported by NIH grant T32NS007222. P.G.P. was supported by NSF grant 1926576, A. Alfred Taubman Medical Research Institute, and NIH grant R01GM111293. C.A.C. was supported by NSF grant 1926576, Craig H. Neilsen Foundation project 315108, NIH grant R01GM111293, and MCubed project 1482.

## AUTHOR CONTRIBUTIONS

M.S.W., P.G.P., and C.A.C. supervised this work and conducted non-human primate surgeries. M.J.M. and A.K.V. wrote and validated software and assisted with conducting experiments. S.R.N. programmed and executed all decoding experiments and data analysis and wrote the manuscript. All authors reviewed and modified the manuscript.

