## Supplementary Figures for "Real-Time Linear Prediction of Simultaneous and Independent Movements of Two Finger Groups Using an Intracortical Brain-Machine Interface"

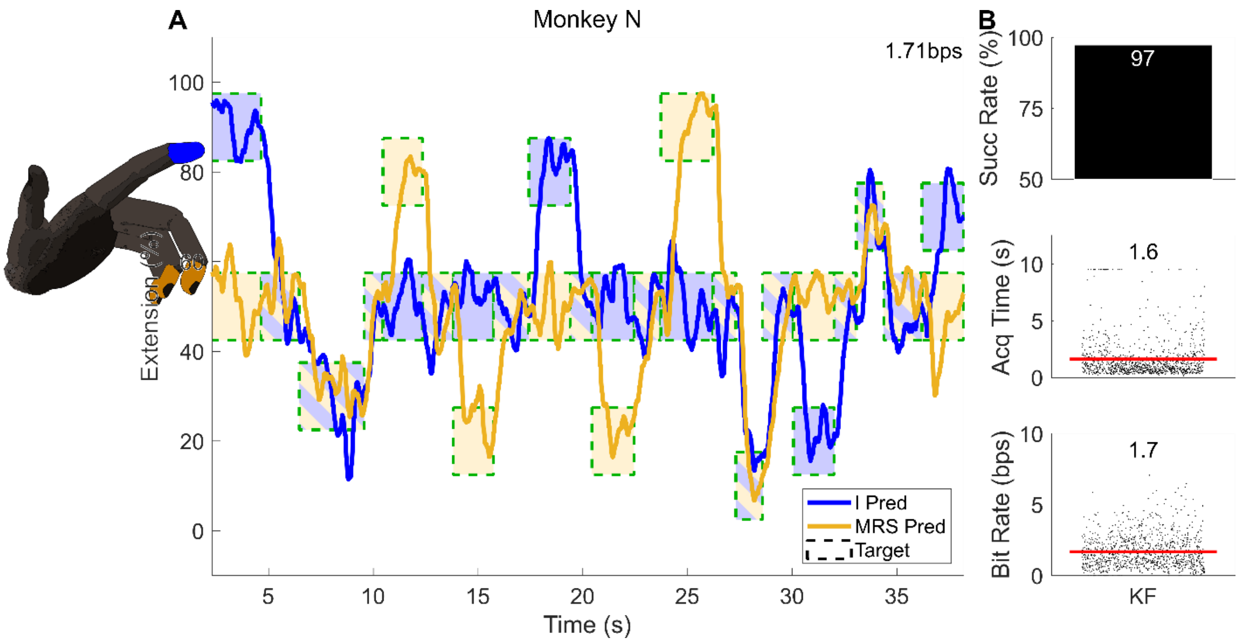


Supplementary Figure 1. Two-finger closed-loop Kalman filter decode using threshold crossing rates. (**A**) Example closed-loop prediction traces from monkey N using the standard Kalman filter, respectively. Targets are represented by the dashed boxes, internally colored to indicate the targeted finger with a border color representing whether the trial was acquired successfully. “I” means the index finger group and “MRS” means the middle/ring/small finger group. The mean bit rate of the trials displayed is presented at the top right. (**B**) Statistics for all closed-loop two-finger threshold crossing rate Kalman filter decodes. The red lines indicate the means, which are numerically displayed above each set of data. The statistic for each trial is represented by one dot in each plot. “Succ Rate” means the percentage of total trials that were successfully acquired in time, and “Acq Time” means target acquisition time.

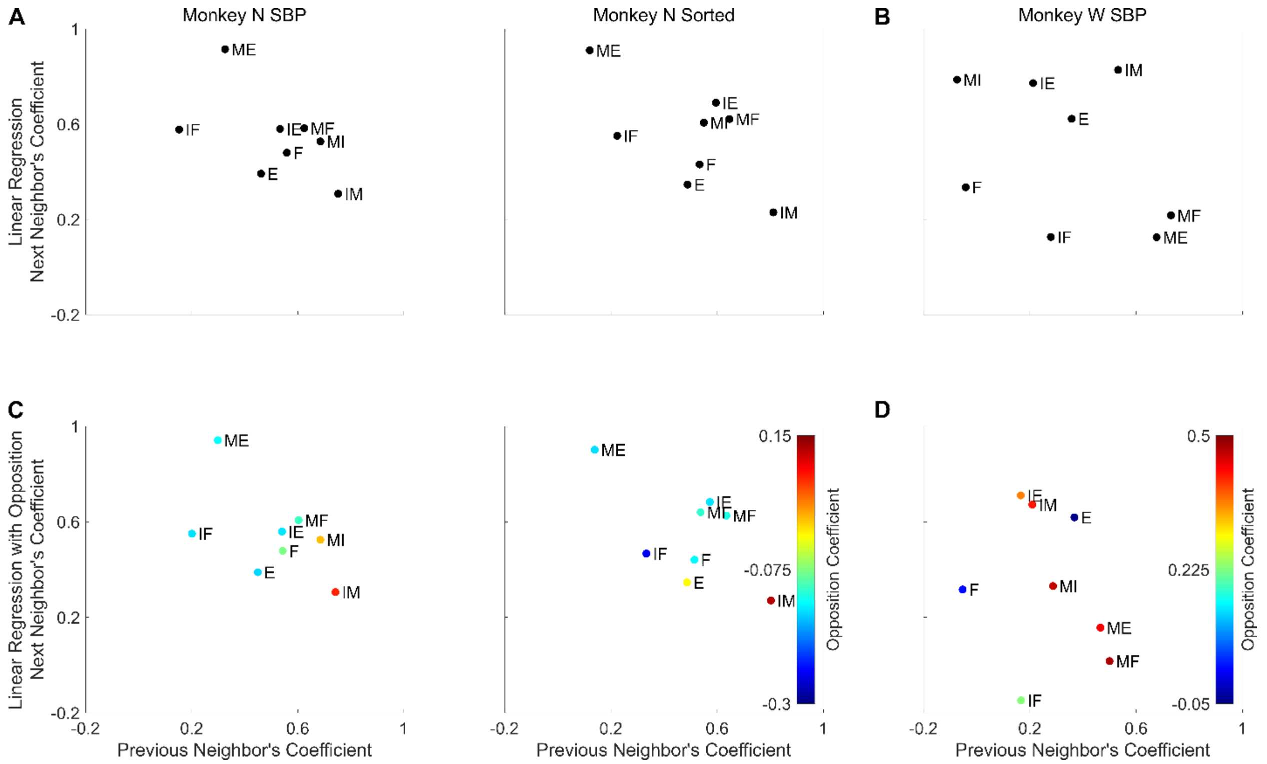


Supplementary Figure 2. Regression coefficients to predict the activity associated with certain movements from the activity of others. Each point is labelled with the movement whose activity would be predicted by the activity of other movements multiplied by the indicated coefficients. I – index finger group, M – MRS finger group, F – flexion, E – extension, IM – index flexion with MRS extension, MI – MRS flexion with index extension. (**A**, **B**) Coefficients for regressions under the assumption that the activity of a movement is a weighted sum of the activities of its neighbors for monkeys N and W, respectively. (**C**, **D**) Coefficients for regressions under the assumption that the activity of a movement is a weighted sum of the activities of its neighbors and the opposite movement for monkeys N and W, respectively. Points are colored based on the best-fit coefficient by which to multiply the opposite movement.

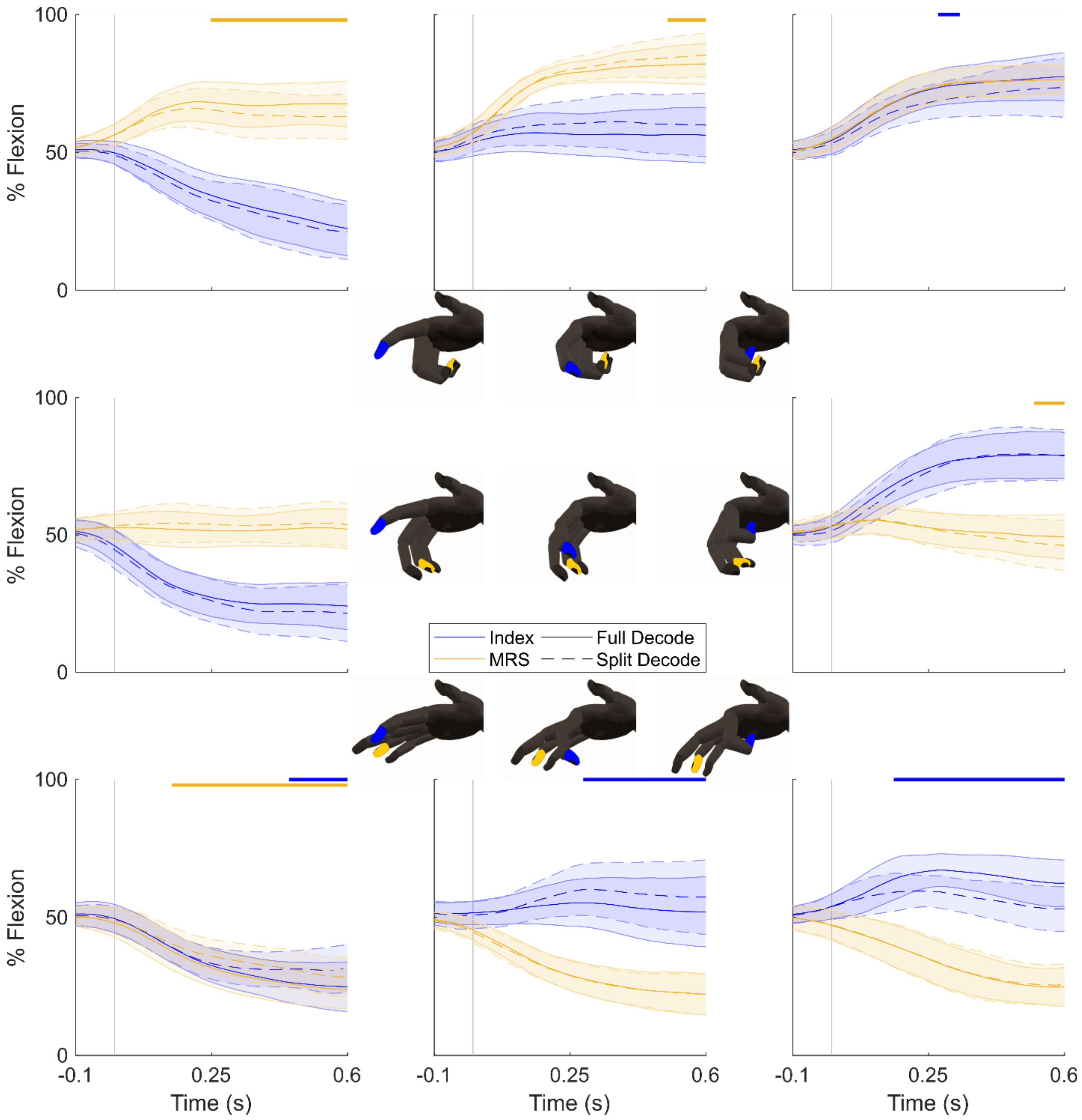


Supplementary Figure 3. Offline ridge regression decoding of all of monkey N's SBP channels, trained on either single or double finger group movements. Single finger group movements (the four middle plots) were decoded using a regression model trained on double finger group movements (the four corner plots), and vice versa for the double finger group movements. These are represented by the "Split Decode" dashed traces. The "Full Decode" solid line traces represent the average decode given the full dataset to train the regression model, with cross-validation. The blue traces correspond to the index group and the yellow traces correspond to the MRS group. The yellow or blue lines near the top of each plot indicate significant differences between the two predicted positions based on a bootstrap analysis on the differences (greater than a one-sided 95% confidence interval). The cartoon hands near each plot demonstrate the movement being decoded, and the letter above each plot indicates which monkey's decodes are displayed.

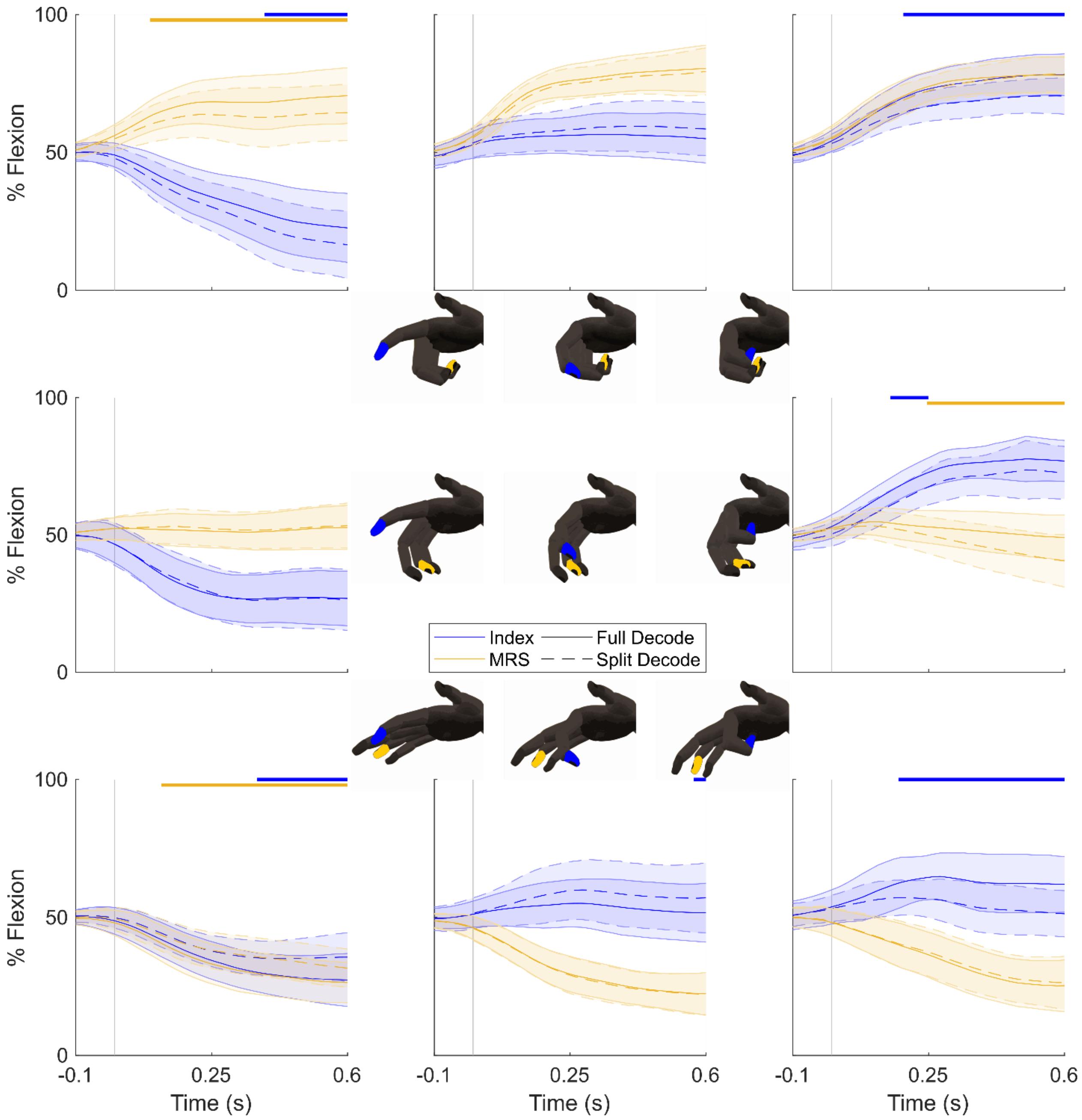


Supplementary Figure 4. Offline ridge regression decoding of all of monkey N's sorted units, trained on either single or double finger group movements. Single finger group movements (the four middle plots) were decoded using a regression model trained on double finger group movements (the four corner plots), and vice versa for the double finger group movements. These are represented by the "Split Decode" dashed traces. The "Full Decode" solid line traces represent the average decode given the full dataset to train the regression model, with cross-validation. The blue traces correspond to the index group and the yellow traces correspond to the MRS group. The yellow or blue lines near the top of each plot indicate significant differences between the two predicted positions based on a bootstrap analysis on the differences (greater than a one-sided 95% confidence interval). The cartoon hands near each plot demonstrate the movement being decoded, and the letter above each plot indicates which monkey's decodes are displayed.

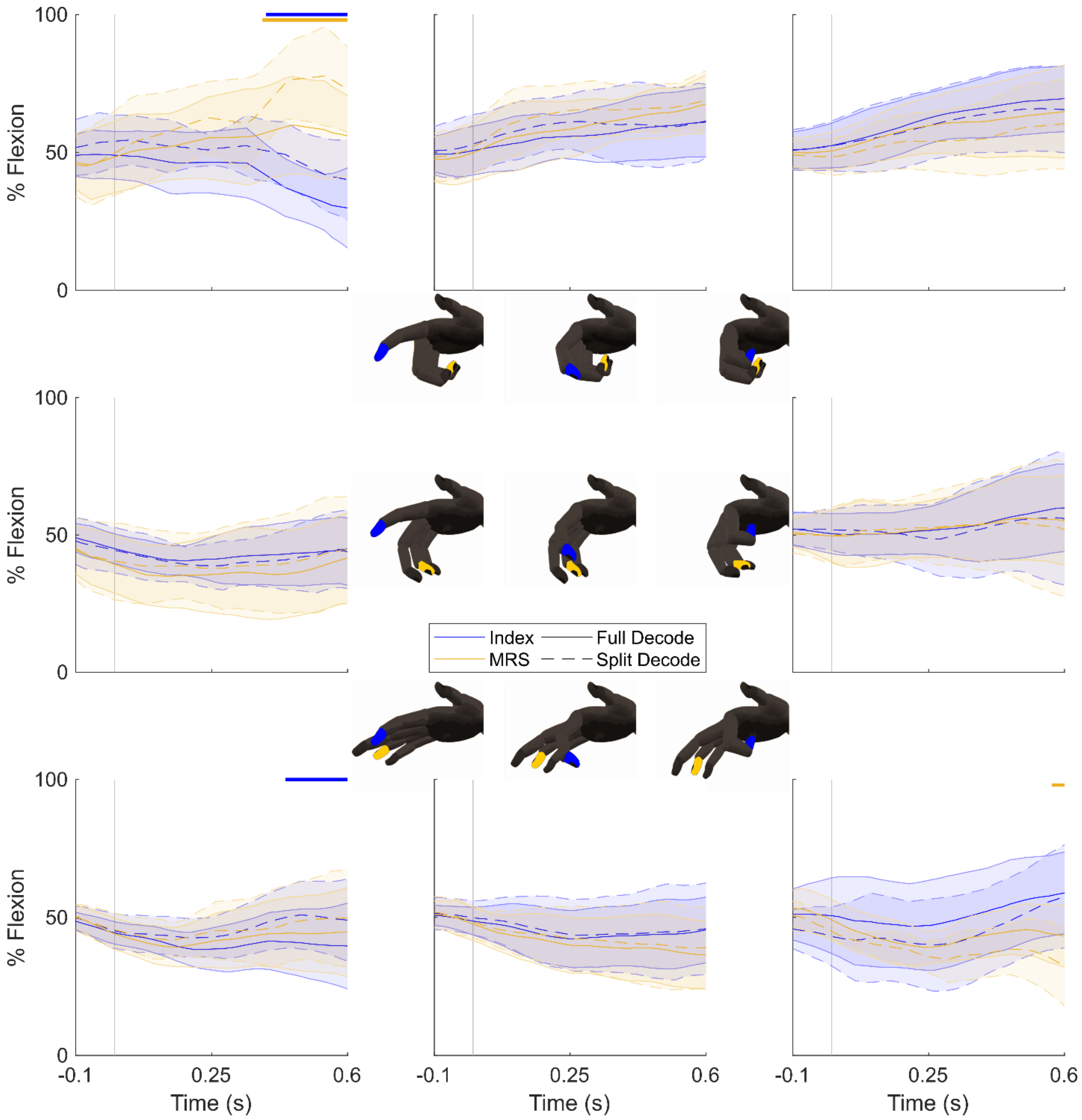


Supplementary Figure 5. Offline ridge regression decoding of all of monkey W's SBP channels, trained on either single or double finger group movements. Single finger group movements (the four middle plots) were decoded using a regression model trained on double finger group movements (the four corner plots), and vice versa for the double finger group movements. These are represented by the "Split Decode" dashed traces. The "Full Decode" solid line traces represent the average decode given the full dataset to train the regression model, with cross-validation. The blue traces correspond to the index group and the yellow traces correspond to the MRS group. The yellow or blue lines near the top of each plot indicate significant differences between the two predicted positions based on a bootstrap analysis on the differences (greater than a one-sided 95% confidence interval). The cartoon hands near each plot demonstrate the movement being decoded, and the letter above each plot indicates which monkey's decodes are displayed.

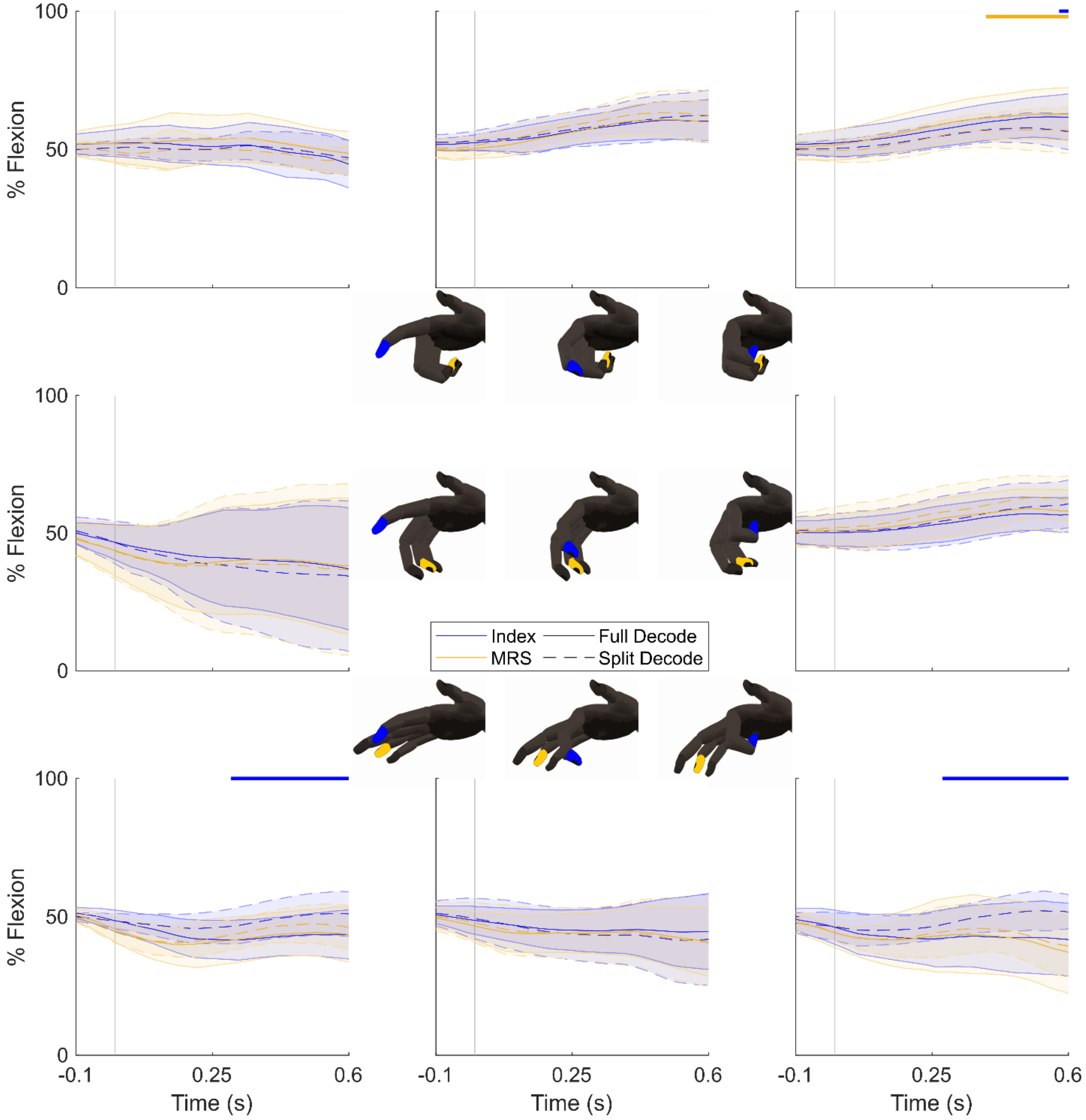


Supplementary Figure 6. Offline ridge regression decoding of all of monkey W's sorted units, trained on either single or double finger group movements. Single finger group movements (the four middle plots) were decoded using a regression model trained on double finger group movements (the four corner plots), and vice versa for the double finger group movements. These are represented by the "Split Decode" dashed traces. The "Full Decode" solid line traces represent the average decode given the full dataset to train the regression model, with cross-validation. The blue traces correspond to the index group and the yellow traces correspond to the MRS group. The yellow or blue lines near the top of each plot indicate significant differences between the two predicted positions based on a bootstrap analysis on the differences (greater than a one-sided 95% confidence interval). The cartoon hands near each plot demonstrate the movement being decoded, and the letter above each plot indicates which monkey's decodes are displayed.

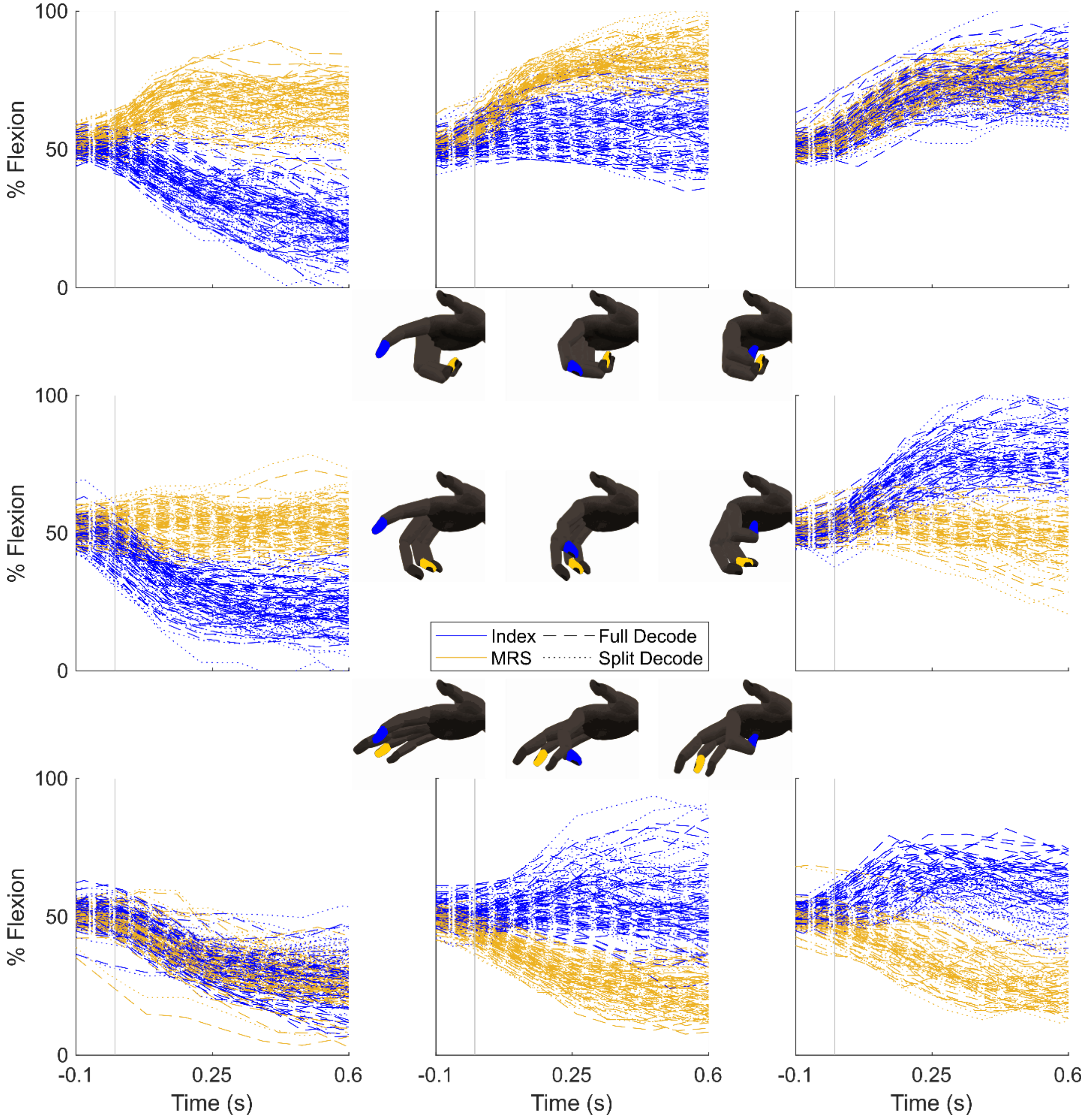


Supplementary Figure 7. Individual trial offline ridge regression decoding of all of monkey N's SBP channels, trained on either single or double finger group movements. Single finger group movements (the four middle plots) were decoded using a regression model trained on double finger group movements (the four corner plots), and vice versa for the double finger group movements. These are represented by the "Split Decode" dotted traces. The "Full Decode" dashed traces represent the average decode given the full dataset to train the regression model, with cross-validation. The blue traces correspond to the index group and the yellow traces correspond to the MRS group. The cartoon hands near each plot demonstrate the movement being decoded, and the letter above each plot indicates which monkey's decodes are displayed.

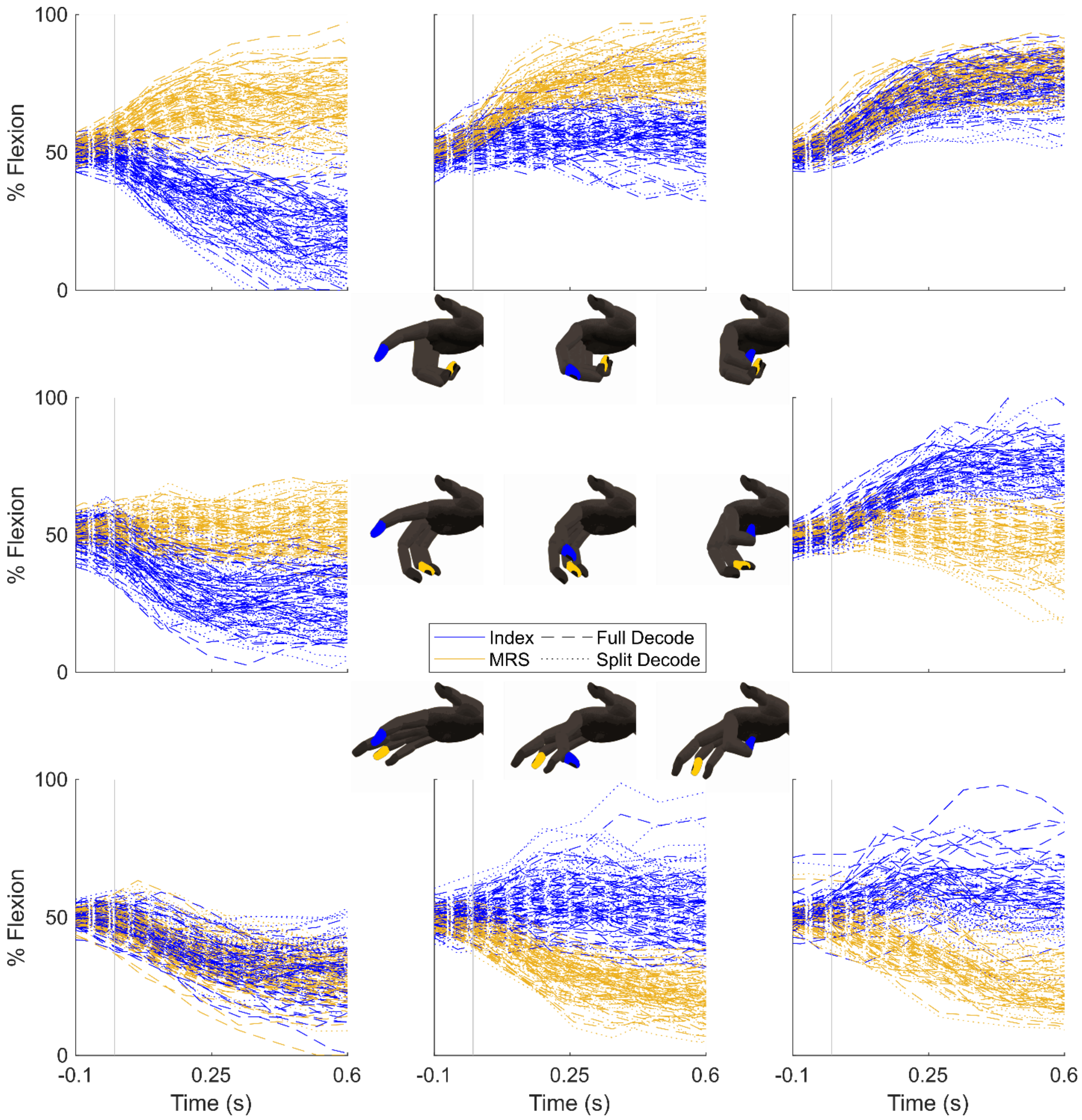


Supplementary Figure 8. Individual trial offline ridge regression decoding of all of monkey N's sorted units, trained on either single or double finger group movements. Single finger group movements (the four middle plots) were decoded using a regression model trained on double finger group movements (the four corner plots), and vice versa for the double finger group movements. These are represented by the "Split Decode" dotted traces. The "Full Decode" dashed traces represent the average decode given the full dataset to train the regression model, with cross-validation. The blue traces correspond to the index group and the yellow traces correspond to the MRS group. The cartoon hands near each plot demonstrate the movement being decoded, and the letter above each plot indicates which monkey's decodes are displayed.

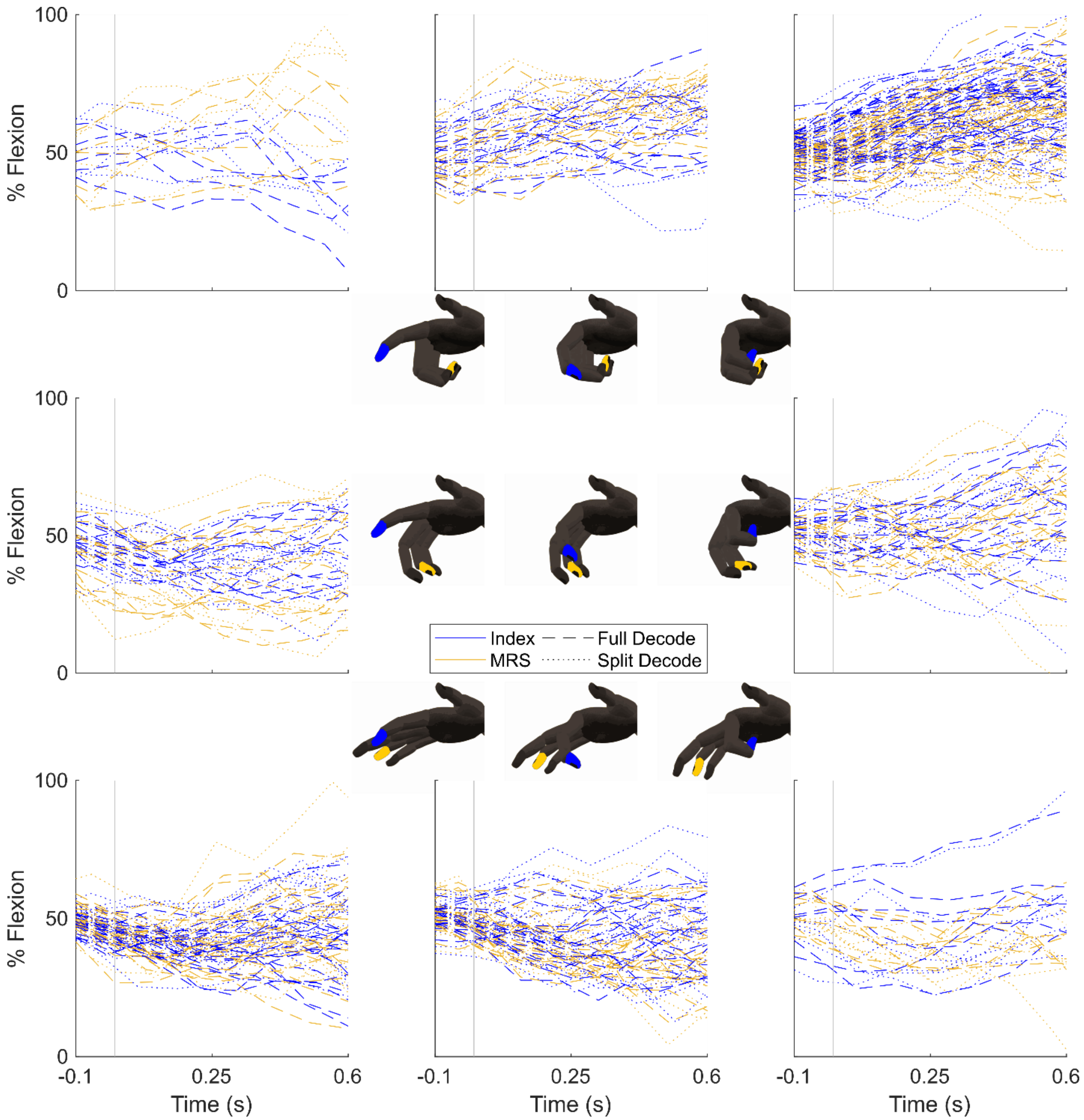


Supplementary Figure 9. Individual trial offline ridge regression decoding of all of monkey W's SBP channels, trained on either single or double finger group movements. Single finger group movements (the four middle plots) were decoded using a regression model trained on double finger group movements (the four corner plots), and vice versa for the double finger group movements. These are represented by the "Split Decode" dotted traces. The "Full Decode" dashed traces represent the average decode given the full dataset to train the regression model, with cross-validation. The blue traces correspond to the index group and the yellow traces correspond to the MRS group. The cartoon hands near each plot demonstrate the movement being decoded, and the letter above each plot indicates which monkey's decodes are displayed.

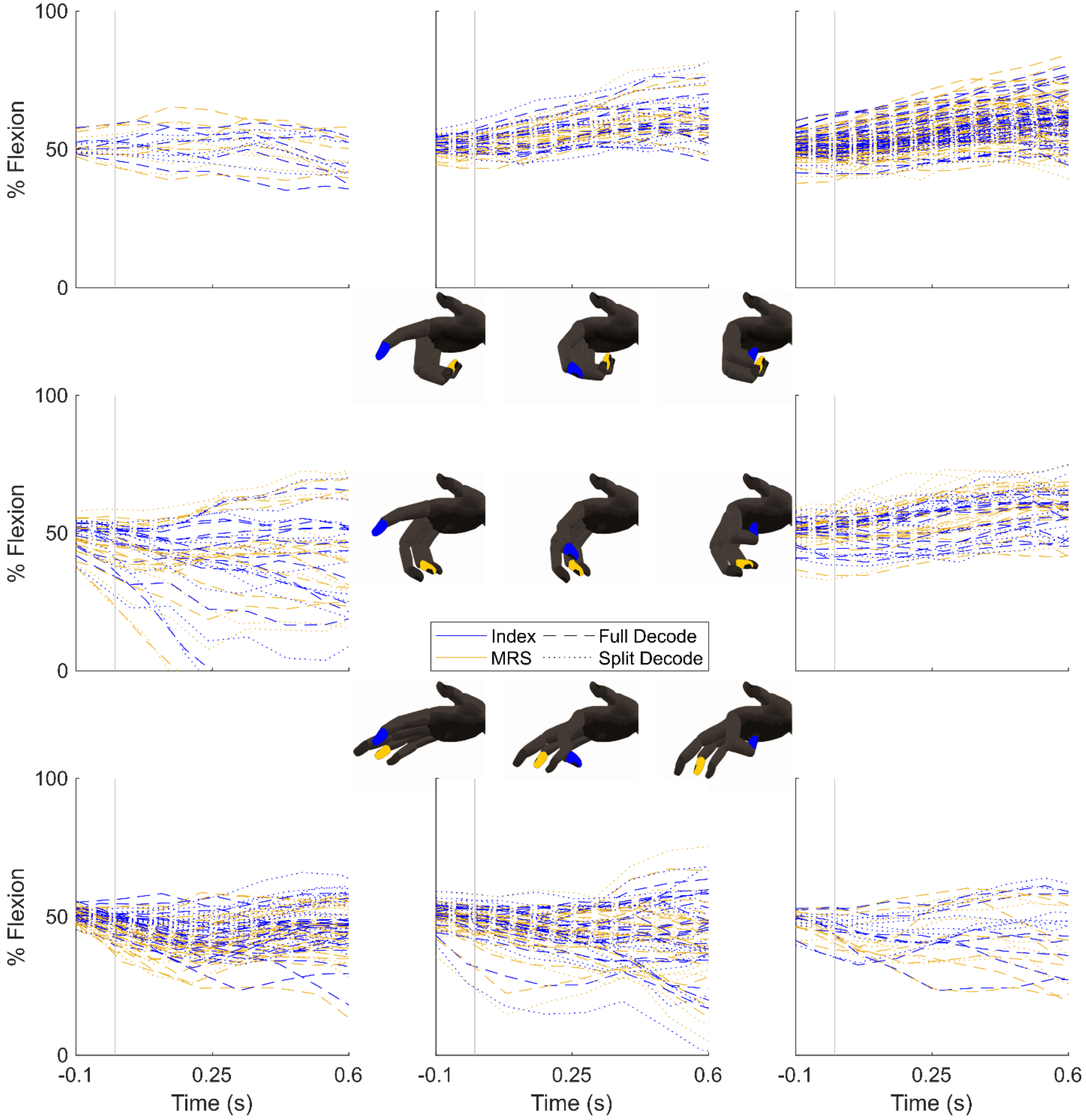


Supplementary Figure 10. Individual trial offline ridge regression decoding of all of monkey W's sorted units, trained on either single or double finger group movements. Single finger group movements (the four middle plots) were decoded using a regression model trained on double finger group movements (the four corner plots), and vice versa for the double finger group movements. These are represented by the "Split Decode" dotted traces. The "Full Decode" dashed traces represent the average decode given the full dataset to train the regression model, with cross-validation. The blue traces correspond to the index group and the yellow traces correspond to the MRS group. The cartoon hands near each plot demonstrate the movement being decoded, and the letter above each plot indicates which monkey's decodes are displayed.

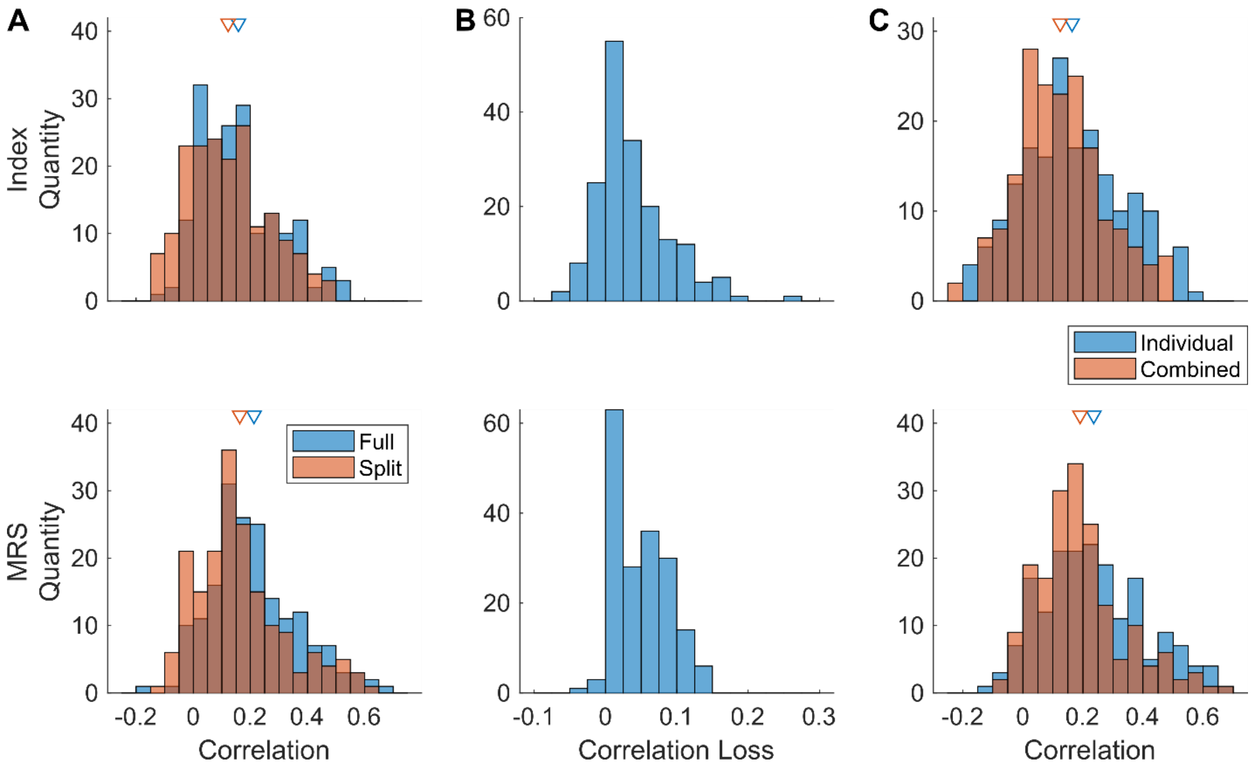


Supplementary Figure 11. Finger-tuned neural activity generalization to untrained behaviors. One datapoint is either one sorted unit from monkey N or one SBP channel from monkey N or monkey W (total 171). Arrows indicate the means. (**A**) Histogram of the Pearson's correlation coefficients between offline predicted movements and true behavior. In orange are the correlations from regressions trained on either single or double finger movements and tested on the opposite, and in blue are the correlations from regressions trained and tested with cross-validation on the full dataset. (**B**) Histogram of the loss in Pearson's correlation between the full dataset training and testing and the training and testing of the dataset split by number of finger groups. (**C**) Histogram of the Pearson's correlation coefficients for the regressions trained on single group movements (blue, tested on double group movements) and the regressions trained on double group movements (orange, tested on single group movements).

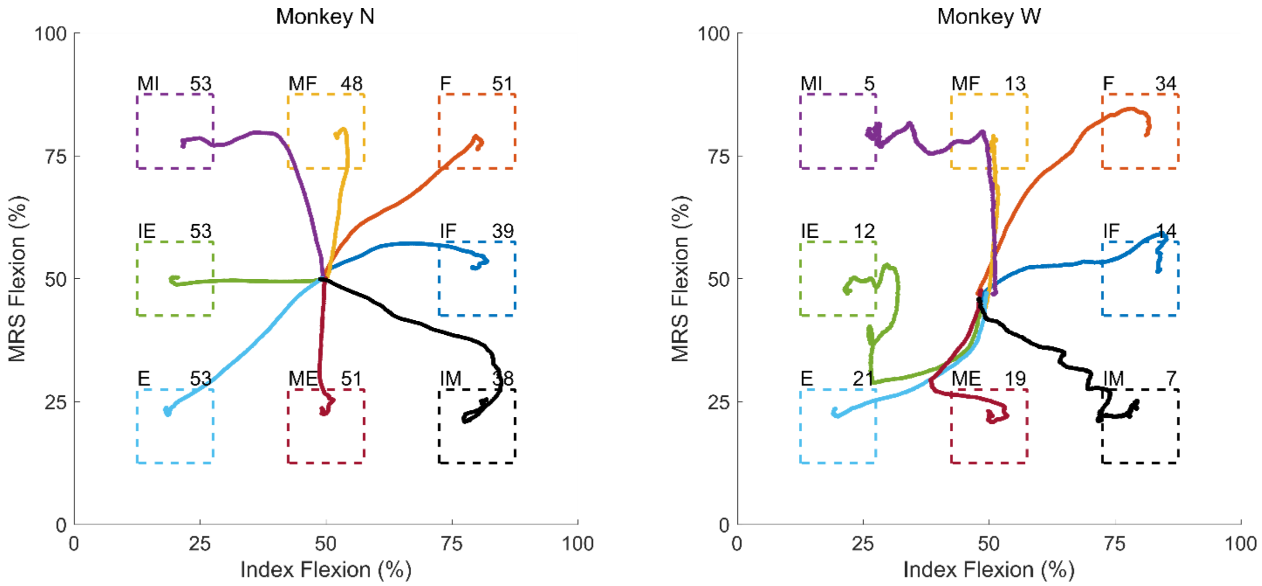


Supplementary Figure 12. Averaged example behaviors for each monkey (N left, W right). Each two-dimensional trace is plotted as MRS flexion percentage versus index flexion percentage, in the same space as Figure 1B. Targets are the dashed boxes, color coordinated to the traces corresponding to attempts to acquire that target. The letters to the top left of each target detail the movement type in the two-dimensional space, and the numbers to the top right indicate the number of trials used to obtain each averaged trace. The behaviors to generate this figure were from the same data sets upon which the tuning analyses were performed. Note that most trajectories are directed towards the targets as if this were one two-dimensional task, but the IM and MI trajectories for both monkeys and the IE and ME trajectories for monkey W suggest the monkeys may have viewed the task as two one-dimensional tasks.
